# Structure-informed mutagenesis identifies a conserved region critical for mouse insulin receptor 5’UTR IRES function

**DOI:** 10.1101/2024.11.05.622145

**Authors:** William B. Dahl, Tammy C.T. Lan, Silvia Rouskin, Michael T. Marr

## Abstract

Cells under stress shift their proteome by repressing cap-dependent translation initiation. RNA elements called internal ribosome entry sites (IRES) can allow key cellular transcripts to remain efficiently translated to support an effective stress response. Well-characterized IRESes depend on RNA structures that reduce the protein requirements for translation initiation, thus circumventing translation inhibition. We have previously determined that the insulin receptor 5’ untranslated region (5’UTR) possesses a capacity for IRES activity that is conserved from insects to mammals. There are several prominent examples of viral IRES structures solved in solution; however, the RNA secondary structures of cellular IRESes remain mostly elusive, especially *in vivo*. Here we probe the secondary structure of the Insr 5’UTR IRES in tandem with two well-studied viral IRESes from Hepatitis C virus (HCV) and Encephalomyocarditis virus (EMCV) using dimethyl sulfate mutational profiling by sequencing (DMS-MaPseq) in cells and *in vitro*. We find that the viral IRES structures in cells are consistent with their known *in vitro* structures and that significant linearization of these well-studied IRESes occurs in the region surrounding their translation start codon in cells. Using the concurrent DMS-MaPseq probing as a constraint, we present a model of the mouse insulin receptor (Insr) 5’UTR. With this model as a guide, we employed a mutation strategy which allowed us to identify a conserved segment distal from the translation start codon as critical for Insr IRES function. This knowledge informed the design of a minimal IRES element with equivalent activity to the full-length Insr 5’UTR across translation contexts.

**Background:** The RNA structural requirements for cap-independent translation initiation facilitated by the *Mus musculus* insulin receptor (Insr) 5’ untranslated region (5’UTR) are unknown.

**Results:** RNA secondary structure probing of the Insr 5’UTR in cells provides a folding model used to identify elements required for cap-independent translation initiation.

**Conclusion:** A small region of the Insr 5’UTR distal to the translation start codon is necessary for cap-independent translation initiation and is fully sufficient in the proper structural context.

## INTRODUCTION

Cells under stress modify their translation environments, sacrificing protein synthesis to avert the accumulation of deleterious products and to divert energy to stress response pathways. One means of translation suppression is the dephosphorylation of the inhibitory eukaryotic translation initiation factor 4E binding protein (4E-BP). Hypophosphorylated 4E-BP binds to eIF4E and impairs formation of the eIF4F cap-binding complex, inhibiting cap-dependent translation initiation^1^. When cap-mediated recruitment of mRNAs to ribosomes is disrupted, an estimated 3-5% of specific cellular transcripts remain efficiently translated^2,3,4,5^. Cap-independent translation initiation is key for mounting an appropriate response to stressors, and this capability can be mediated by RNA structures called internal ribosome entry sites (IRES) in mRNA 5’ untranslated regions (5’UTRs)^6^. Because structured elements in the 5’UTRs of model mRNAs inhibit cap-dependent translation initiation^7,8^, efficient translation of mRNAs with highly structured 5’UTRs may rely on IRESes or other IRES-like translation enhancer sequences^9^.

IRES activity was first discovered and is well-studied in RNA viruses^10,11^. The RNA structures and mechanisms of several viral IRESes are known^12,13,14,15^. Viral IRESes tend to have compact, highly conserved RNA structures with potent IRES activity. This may reflect the fitness objective to rapidly produce proteins necessary for viral proliferation^16^. By contrast, cellular IRESes present cells with an opportunity to specifically regulate protein synthesis^6^, and IRES elements in cellular transcripts may have less stringent structural requirements^9,17^. Additionally, endogenous cellular transcripts may be translated by both cap-independent and cap-dependent mechanisms. Classification of structural motifs and IRES mechanisms in endogenous cellular transcripts remains elusive^18^.

Previous work in our lab found that the 5’UTRs of insulin-like receptors possess a capacity for cap-independent translation initiation which is conserved from flies to mammals^19,20^, despite a great divergence in sequence. Insulin receptor mRNA is transcriptionally upregulated by fasting conditions in various organisms^21,22,23^. Transcriptional control exerted on insulin receptor directly parallels its protein expression in insects and mammals^24^, despite the repression of cap-dependent translation in starving cells^25^. It is thought that the 5’UTR IRES in insulin-like receptors couples transcript expression to protein expression to ensure that the cell rapidly returns to a healthy state when insulin-like signals are received. Known viral IRES mechanisms depend directly on RNA structure, but nearly all published structures have only been probed *in vitro*. The structure of the mouse Insr 5’UTR has not been probed in any context. The cellular environment constrains the range of allowed RNA structures compared to RNA in solution^26^, and probing in cells may provide an opportunity to find the pairing state responsible for IRES activity.

To investigate the RNA secondary structure of the murine Insr 5’UTR, we employed dimethyl sulfate mutational profiling by sequencing (DMS-MaPseq) in cells and *in vitro*. Dimethyl sulfate (DMS) has long served as a chemical probing reagent for RNA structure^27^ because it specifically methylates cytosines and adenosines on their base-pairing faces (N^3^C and N^1^A)^28^. Paired bases are shielded from modification^29^. In DMS-MaPseq, these events are encoded in cDNA synthesis as mutations, which is more conducive to probing large RNA structures than truncation-based methods^30^. To validate our approach, we probed two viral IRESes with well-characterized structures from the 5’UTRs of Encephalomyocarditis virus (EMCV) and Hepatitis C virus (HCV). While the RNA genome of HCV has been probed in cells^31^, the EMCV viral IRES has only been probed *in vitro*. We used these data to constrain RNA secondary structure predictions, allowing us to model the base-pairing state of the Insr 5’UTR.

Informed by this model, we generated a series of mutations aimed at elucidating Insr IRES function. To investigate whether structure plays a significant role in the Insr 5’UTR’s IRES activity, we also probed key mutants with DMS-MaPseq in cells and *in vitro*. We identified the necessary and sufficient portions of the UTR for IRES activity by assaying the cap-independent performance of the engineered RNA secondary structures. Our results suggest that the conserved elements distal to the translation start codon are critical and sufficient, but that a sequence- and structure-independent aspect of the remaining UTR is necessary. This minimal sufficient segment has equivalent cap-independent performance to the full-length UTR in cells and *in vitro*. This work provides a model of the RNA secondary structure of the *Mus musculus* Insr 5’UTR in cells and uses this model to determine which portion contributes to IRES activity, a key step to understanding the nature of cellular IRESes.

## MATERIALS AND METHODS

### DMS treatment and RNA extraction

HEK293T cells (from ATCC, Cellosaurus RRID: CVCL_0063) were maintained in Dulbecco’s Modified Eagle’s Medium (Genesee Scientific), with added 10% Fetal Bovine Serum (Gemini Bio), 1µg/mL insulin solution from bovine pancreas (MilliporeSigma), and 1X penicillin/streptomycin (Fisher Scientific) at a subculture ratio of 1:6. For transfections, 10cm tissue culture plates were seeded with 3x10^6^ cells each. After 24 hours, the cells were cotransfected with a *Gaussia* luciferase reporter containing HCV, EMCV, and Insr IRESes in their 5’UTRs using 48µL 1mg/mL polyethylenimine (PEI) and 4µg of each plasmid (a 4:1 µL PEI to µg DNA ratio). For mutants of Insr, 12µg of plasmid were transfected. After 48h, the cells reached a confluency of ∼80% and were treated with dimethyl sulfate (DMS). 200µL (2%) DMS was dispersed dropwise into the DMEM atop the cells in 10cm plates. The plates were incubated at 37°C for 5 minutes, and the reaction was quenched by adding 10mL 30% beta-mercaptoethanol. RNA was extracted by scraping the cells and harvesting in 15mL conical tubes. The cell pellet was homogenized in 1mL TRI-Reagent (Molecular Research Center, Inc.), and 200µL chloroform was pipette mixed, incubated for 1 minute at room temperature, and then centrifuged at 21,000 x g for 15 minutes at 4°C. The aqueous phase was transferred to a new tube and mixed with one volume of 100% ethanol. RNA was then purified using the Zymo Clean and Concentrator-25 kit following the manufacturer’s protocol with DNaseI digestion in the column.

### DMS probing *in vitro*

*In vitro* transcribed RNA containing HCV, EMCV, Insr, and Igf1r UTR sequences were pooled. *In vitro* probing was conducted as described.^1^ Briefly, 1µg of *in vitro* transcribed RNA containing the structures of interest was added to a mixture containing 26µg of calf liver tRNA to a total volume of 100µL. The tRNA was added to ensure identical RNA:DMS ratios between these probing experiments and subsequent probing of single mutants. These RNAs were refolded by denaturing at 95°C for 1 minute, and then placing them on ice. Sodium cacodylate (final concentration 100mM) and MgCl2 (final concentration 6mM) were added to a yield a final volume of 292.5µL. The RNA was incubated at 37°C for 5 minutes to reach folding equilibrium^2^. 7.5µL of DMS (final [DMS] 2.5%) was added and the probing reaction was mixed at 37°C, 500rpm on a thermomixer with a 1.5mL tube attachment for 5 minutes. The reaction was neutralized by adding 150µL of 100% beta-mercaptoethanol. The probed RNAs were then cleaned using the recommended RNA Clean and Concentrator-5 protocol (Zymo).

### Reverse transcription for mutational profiling

Reverse transcription (RT) was performed with maltose binding protein (MBP) fused TGIRT-III purified with amylose resin as previously described^32^. All DNA oligos were purchased from Genewiz, Inc., and their sequences are listed in Table 1. 2pmol of RT primer specific to our targets was annealed to the RNA by incubating at 75°C for 10 minutes in a thermocycler and ramping down 0.2°C/s to 25°C. After reaching 25°C, a master mix containing RT buffer (final concentration: 10mM NaCl, 10mM MgCl2, 20mM Tris-HCl pH 7.5, 1mM DTT), 1µL TGIRT-III, and 10mM dNTPs (BioBasic) was added and mixed. The RT reaction was incubated at 25°C for 30 minutes and then 60°C for 90 minutes, followed by 10 minutes at 70°C. 1µL (5U) of RNaseH (NEB) was added and the reaction was incubated for 30 minutes at 37°C. The cDNA was then purified using DNA cleanup columns (BioBasic).

### Target-specific DMS-MaPseq library preparation

Polymerase chain reaction was performed on the cDNA targeting tiled amplicons along each 5’UTR for a maximum of 27 cycles using PfuX7 DNA polymerase. His-tagged PfuX7 was expressed from pET-PfuX7 in BL21*(DE3) *E. coli* cells (ThermoFisher) containing the pLacIRARE2 plasmid (Novagen) and purified with nickel resin as previously described^33^. The primers for these reactions contained Illumina adaptor sequence as overhangs. For forward primers, this overhang was: 5’- ACACGACGCTCTTCCGATCT-3’; reverse primers: 5’-AGACGTGTGCTCTTCCGATCT-3’. These fragments were purified using a column cleanup to remove unused oligos (BioBasic). A subsequent PCR of 6-8 cycles was used to add P5/P7 attachment sequences and a single barcode to 300ng of the initial PCR product for a completed library, and these amplicons were column purified again. Up to 70 library fragments were pooled into each sequencing run on a MiSeq Micro PE150 flow cell (Illumina). To sequence multiple replicates and fragments with common primers, the resultant FASTQ files were barcode split using Barcode Splitter hosted on Galaxy^34^. These barcodes were the first 6bp of each target-specific primer, which are the first bases in each read. The split reads were interlaced using FASTQ interlacer to separate fragments with only one unique primer, and then deinterlaced using FASTQ de-interlacer to differentiate fragments^35^. Interlacing and de-interlacing was also conducted on Galaxy^34^. The data were analyzed using the established Detection of RNA folding Ensembles by Expectation Maximization (DREEM) pipeline, where mapped reads containing mutations are converted to bit vectors as described^36^. Data was filtered to exclude reads containing mutations spaced less than 4nt apart, reads with a proportion of mutations exceeding 10% of their length, and reads with a quantity of mutations exceeding the median number of mutations by 3 standard deviations as previously described^36,37^. In these experiments, frequency histograms tracking mutations were typically centered on 2 mutations/read (Supp Fig. 1d-e, g-h, and k-l), which corresponds to previous work titrating DMS treatments in various cellular contexts^26,30,36^.

**Figure 1.**
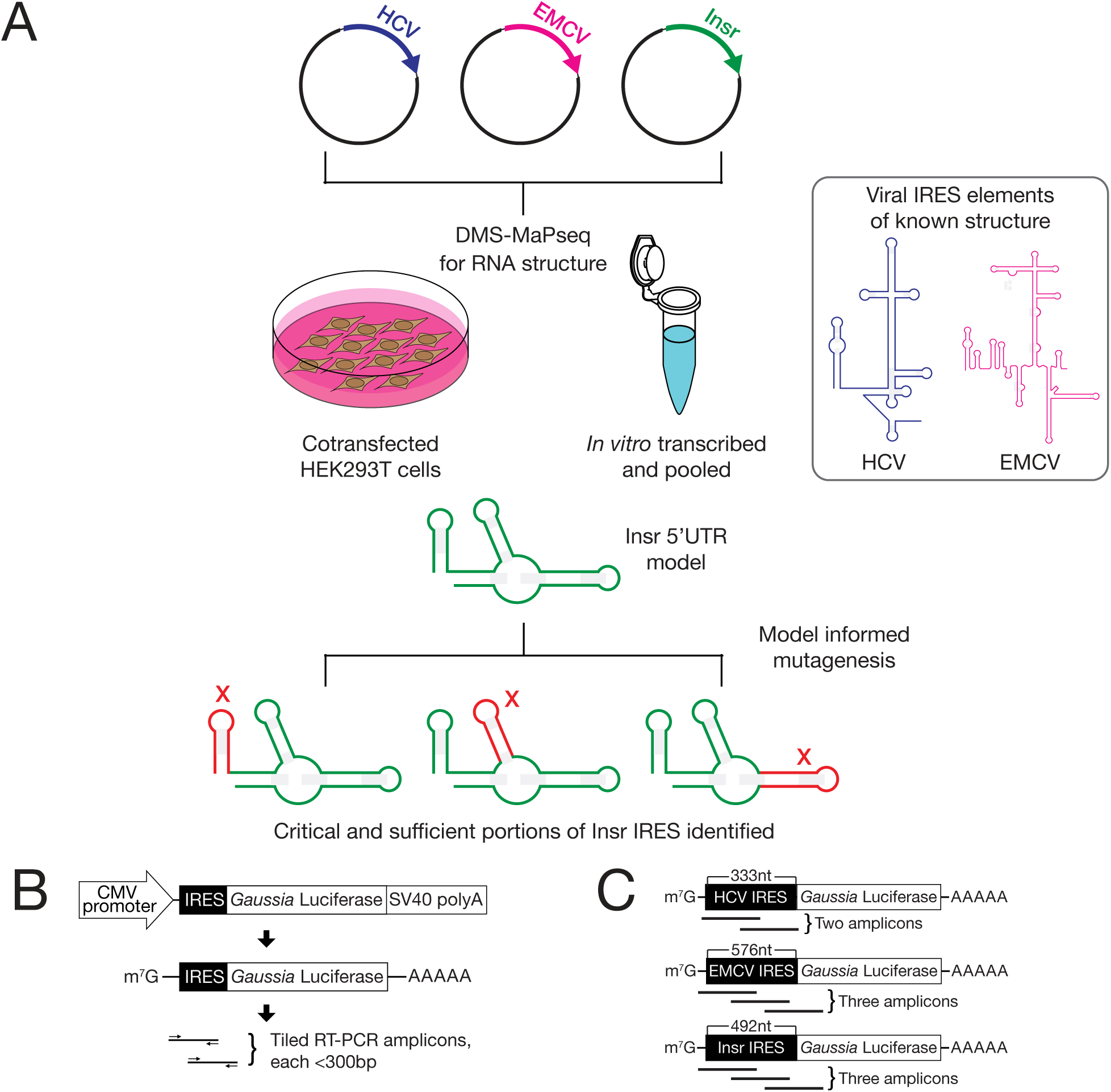
DMS-MaPseq in cells constrains RNA secondary structure predictions of internal ribosome entry sites (IRESes) **(A)** Three luciferase reporter plasmids containing IRES elements in their 5’ untranslated regions (5’UTRs) were cotransfected into HEK293T cells. Two viral IRESes from hepatitis C virus (HCV) and encephalomyocarditis virus (EMCV) 5’UTRs were probed with dimethyl sulfate (DMS) alongside the mouse insulin receptor (Insr) 5’UTR. Resultant transcripts are depicted with tiled RT-PCR amplicons used to prepare DMS-MaPseq libraries using the established gene specific strategy. These RNA structures were also pooled and probed *in vitro*. Prediction of the Insr 5’UTR RNA secondary structure informed a mutation strategy aimed at identifying functional components. **(B)** Diagram of plasmid expression cassette for this strategy. **(C)** Diagram of transcripts probed, with number of tiled RT-PCR amplicons annotated for each 5’UTR.

### DMS-constrained RNA secondary structure prediction

Secondary structure modeling was conducted using RNAstructure via the RNAstructure Python Interface^38^. Three input metrics were set for the predictions: normalized bases, which specifies the mutation frequency corresponding to 1.0 in DMS signal normalization; maximum pairing distance, which restricts long distance basepairing; and signal threshold, under which all mutation frequencies are set to 0. Each base’s normalized DMS reactivity is on a scale of 0 to 1 and is expressed as a proportion of the most reactive set specified in normalized bases. Bases with normalized DMS reactivity of 0-0.4 are colored blue in all plots, 0.4-0.8 yellow, and 0.8-1.0 red. For all models reported, normalized bases are 5% the length of the predicted structure rounded to the nearest whole number, maximum pairing distance is 250nt, and the signal threshold is a 0.005% mismatch rate. Prediction used an energy function previously described for DMS and SHAPE mutational profiling which does not use an empirically determined slope and intercept^39^.

### ΔDMS calculation and arc plots

Differences in DMS reactivities were calculated by first normalizing the mutation rates using the same method as described: setting the median mutation frequency of the top 5% of bases equal to 1.0 and expressing all mutation rates as a proportion of this median. Normalized values from *in vitro* probing data were subtracted from normalized in-cell data (in cell – *in vitro*). The associated arc plots represent the same structural predictions constrained by *in vitro* DMS-MaPseq data presented in this report.

### Calculating modified Fowlkes-Mallows Index (mFMI) for structure comparison

The Fowlkes-Mallows Index (FMI) is used as a metric for base-pairing similarity between two RNA secondary structures ranging from 0 (no similarity) to 1 (identical); FMI is calculated as the geometric mean of the true positive rate (TPV) and the positive predictive value (PPV), and it was originally used to compare hierarchical clustering^40^. For RNA structure comparison, TPV is the proportion of identical basepairs out of identical basepairs plus false negatives, and PPV is the proportion of identical basepairs out of identical basepairs plus false positives. The mFMI used in this report factors in shared unpaired bases, and it is calculated as previously described^37^: by multiplying the FMI by (1 – the proportion of shared unpaired bases), which weights the FMI to the proportion of paired bases, and then adding the proportion of shared unpaired bases to get the final mFMI value. With *u* representing the proportion of shared unpaired bases, mFMI = *u* + (1 – *u*)*FMI. mFMI in this report is calculated in sliding windows of 30nt in 1nt increments, plotting the result to the fifteenth nucleotide in the window. mFMI was computed with the DREEM pipeline.

### Calculating the area under the receiver operator characteristic curve (AUROC)

The area under the receiver operator characteristic curve (AUROC) reflects signal-to-structure agreement between the DMS-MaPseq data and the predicted structure. This assumes that the reactivity of unpaired bases should be higher than that of paired bases. The predicted secondary structure is used to classify a given base as paired or unpaired. DMS reactivity values exceeding a sliding threshold and classified as paired are considered false positives, while paired bases below the threshold are considered true positives. The AUROC values were computed with SciPy^41^.

### Site-Directed Mutagenesis

Site-directed mutagenesis was conducted with whole-plasmid PCR, using divergent primers that exclude the deletions of interest and add sequence as overhangs for the specified mutants. All plasmids were sequenced through their luciferase ORFs by Sanger sequencing and plasmids with effects were fully sequenced via whole-plasmid sequencing. For plasmids with multiple mutagenesis sites, consecutive plasmids were generated. A list of all primers is contained in Table 1. A list of all plasmids’ mutagenesis sites and the primers used to generate them can be found in Table 2.

### Bicistronic luciferase assays in cells

Plasmids containing bicistronic luciferase reporters driven by RSV promoter were transfected into HEK293T cells on 24-well tissue culture treated plates 24 hours after seeding 50,000 cells/well. Polyethylenimine (PEI) was used in a ratio of 4µL 1mg/mL PEI for 1µg plasmid, with 600ng of plasmid per well. Three technical replicates were executed per mutant, with beta-globin 5’UTR as a non-IRES control and HCV IRES as a positive control. After 48 hours, the cells were harvested at ∼80% confluency. The DMEM was removed, and the cells were washed once with 200µL 1X PBS rocking for 5 minutes at room temperature. The PBS was removed and 100µL of 1X Passive Lysis Buffer (Promega) was added. The cells were lysed by rocking for 20 minutes at room temperature and then 10µL of lysate was assayed immediately, taking care not to aspirate any cell debris. First, 100µL of firefly luciferase buffer were added and the well was read, then 100µL of Renilla luciferase buffer were added and the well was read again. Firefly luciferase buffer was: 75mM Tris pH 8.0, 5mM MgSO4, 0.1mM EDTA, 0.53mM ATP, 5mM D-luciferin (GoldBio). Renilla luciferase buffer was: 25mM Na4PPi, 10mM NaOAc, 15mM EDTA, 500mM Na2SO4, 1M NaCl, 10mM coelenterazine (GoldBio), adjusted to pH 5.0. The resulting luciferase values were read on a Synergy HTX plate reader (BioTek). Raw luciferase assay data for all experiments can be found in Table 3.

### Bicistronic luciferase assay signal normalization (IRES activity)

To compute IRES activity, the average firefly:Renilla signal in the mutant UTRs was divided by the average firefly:Renilla signals of technical replicates of the full length Insr 5’UTR reporter, multiplying by 100 to produce a percentage. This average is reported for each biological replicate as individual points in the IRES activity plots. Error bars are standard deviation of biological replicates, except for full-length (which is 100% of itself in every assay). The error bars for full-length Insr are standard deviation of technical replicates, averaged across all biological replicates. These computations can be found in Table 3. Statistical comparisons were performed by one-way analysis of variance (ANOVA) followed by Tukey’s honestly significant difference (HSD) test using MATLAB R2022b^42^. Significance indicators are as follows: p ≤ 0.0005 (***), p ≤ 0.005 (**), p ≤ 0.05 (*), p ≥ 0.05 (not significant, ns). Asterisks/ns to the right of each plot are in comparison to full-length UTR, while individual comparisons of interest are denoted in each plot.

### Comparing DMS-MaPseq signal with R^2^ on adenosines and cytosines

To compare in a structure-naïve manner, DMS-MaPseq data from the mutant constructs was directly compared to that of the full-length UTR. In MATLAB R2022b^42^, coefficient of determination (R^2^) was calculated in windows of 31nt plotted from the center, considering only values on As and Cs. Only windows containing positions of shared sequence are plotted. Windows with eight or fewer As and Cs were discarded. The deletions or tetraloop hairpin replacements are depicted in each plot.

### *In vitro* transcription

The plasmids containing the mutant UTRs were used to generate a PCR template for *in vitro* transcription. For *in vitro* translation luciferase assays, the same plasmids used for in-cell luciferase assays were used to create the *in vitro* transcription template. In PCR, forty-three adenines were added by the reverse primer, and a T3 RNA Polymerase promoter was added (Primers 70 and 71). Reporters with different 5’ ends used unique primers ΔD1: (Primer70), BG (Primer71), and HCV (Primer72). For *in vitro* DMS-MaPseq, the same plasmids used for in-cell DMS-MaPseq were used, and *in vitro* transcription templates were prepared by PCR using Primer75 and Primer76. Templates were purified using a column cleanup protocol (BioBasic). All templates were in vitro transcribed using T3 RNA Polymerase (New England Biolabs) in 100µL reactions. The composition of these reactions was: 40mM Tris HCl pH 7.9, 10mM DTT, 2mM Spermidine-HCl, 20mM MgOAc, 7.5mM NTPs (BioBasic), 1U SuperaseIn RNase inhibitor (ThermoFisher), and 0.5U inorganic pyrophosphatase (Sigma). The reactions were incubated for 1 hour at 37°C. The RNA was then cleaned using a Zymo RNA Clean & Concentrator-25 kit according to the manufacturer’s recommendations with the optional in-column DNaseI treatment. The resulting RNA’s purity was assessed by NanoDrop and run on a 1% agarose 1X TBE gel using 2X RNA Loading Dye (New England Biolabs) and stained with 1µg/mL ethidium bromide. The RNA was stored at - 80°C.

### Capping of *in vitro* transcribed RNA

RNAs assayed with *in vitro* translation were capped using the Vaccinia Capping System (NEB). Reactions were conducted according to the manufacturer’s specifications. These reactions were cleaned on Zymo Clean and Concentrator-5 columns. Controls (Insr, BG, and HCV) were capped in three reactions, pooled, and purified with the Zymo Clean and Concentrator-25 kit according to manufacturer recommendations. The resulting elutions were stored at -80°C.

### Monocistronic luciferase assays *in vitro*

Untreated rabbit reticulocyte lysate (RRL) was purchased from GreenHectares, and RRL was treated with hemin by adding 100µL of a 1mM solution to 5mL RRL (20µM final concentration hemin) and mixed immediately upon thawing as previously described^43^. The RRL was not treated with micrococcal nuclease to assay the reporters under competitive conditions. *In vitro* translation assays were conducted in 20µL reactions with 10µL of RRL. The reaction was supplemented with 60µM amino acids, 100ng/µL calf liver tRNA, and creatine kinase energy regeneration mix. Energy generation mix is: 20mM creatine phosphate (Sigma), and 100ng/µL creatine phosphokinase type I from rabbit muscle (Sigma). The buffer was 22mM HEPES pH 7.4 with 0.1mM spermidine, 0.4mM MgOAc, and 30 mM KOAc. Reactions containing 100ng of mRNAs with the cap-dependent beta-globin 5’UTR and Hepatitis C virus IRES were assayed as negative and positive controls, respectively. For comparisons between Insr and mutated reporters, mRNA was matched for molar input: 200ng of Insr RNA (0.27pmol), or 0.27pmol of mutant RNA, varying in mass based on RNA length. The reaction was incubated for 60 minutes at 30°C and stopped by the addition of 1 reaction volume (10µL) of 5X PLB dilute to 2X in water (Promega). Firefly luciferase was then assayed on the Synergy HTX plate reader (BioTek) in the same manner detailed previously. Statistical comparisons follow the same procedure used for the bicistronic luciferase reporters.

### Software used for analysis and for plotting

For analysis: the DREEM pipeline was run locally and uses other programs. DREEM is operated through Python 3.10.15. These analyses used: CutAdapt 3.7 to trim adapter sequences^44^, FastQC 0.12.1 to analyze sequencing quality^45^, and Bowtie2 2.5.4 for alignments^46^. RNAstructure was used for secondary structure modeling^38^. MATLAB R2022b was used for statistics in luciferase assay plots with MATLAB functions anova1 and multcompare^42^. The latest iteration of DREEM pipeline software^36^ can be found at URL: https://codeocean.com/capsule/6175523/tree/v1

For plotting: Python 3.10.15 was used to generate the ΔDMS arc plots with NumPy 1.24.4^47^, pandas 2.2.2^48^, and Matplotlib 3.9.2^49^. VARNA 3.93 was used for RNA secondary structure figures^50^. MATLAB r2022b was used for all other plots^42^. Adobe Illustrator 28.7.1 was used to build the complete figures.

## RESULTS

### The HCV and EMCV IRESes are linearized around their translation start codons in cells

To benchmark DMS-MaPseq for IRES secondary structure prediction, we probed two viral IRESes of known structure (HCV, EMCV) and a cellular IRES of unknown structure (Insr) in tandem (Fig. 1a). HEK293T cells were cotransfected with reporters containing these sequences in the 5’UTRs of a monocistronic *Gaussia* luciferase reporter driven by the Cytomegalovirus (CMV) immediate early promoter (Fig. 1b). We then probed the resultant RNAs with dimethyl sulfate (DMS). This allowed for consistent treatment conditions between HCV, EMCV and the mouse Insr 5’UTR. The HCV, EMCV, and Insr IRESes were also *in vitro* transcribed, pooled, and concurrently treated with DMS *in vitro*. The established gene-specific DMS-MaPseq strategy^30^ was employed to model RNA secondary structure. Libraries were prepared from RT-PCR fragments tiled across each 5’UTR (Fig. 1c). A list of all primers used for this study can be found in Table 1. Each probing experiment was conducted three times unless otherwise specified. The data were analyzed using an established pipeline^36^ with an additional barcode splitting step to support sequencing many samples of similar sequence at once (Supp. Fig. 1a). Elevated mutation rates correspond to a higher likelihood of a base being unpaired^30^, which constrains RNA secondary structure modeling.

The distributions of mutations per sequencing read were observed to ensure a similar frequency of probing events to previous studies both in cells and *in vitro* (Supp. Fig. 1b-m). These DMS probing data were used as constraints for RNA secondary structure prediction in RNAstructure^38^ (see Methods). To gauge the efficacy of our secondary structure modeling, we compared predictions of viral IRES structures to their well-characterized *in vitro* models. The RNA secondary structure models in this report are presented in three contexts: (1) DMS-MaPseq probing in cells, (2) DMS-MaPseq probing *in vitro*, and (3) known structures from previous reports informed by *in vitro* probing. For clarity, we will refer to these three variants as “in-cell”, “*in vitro*”, and “known”, respectively.

Folding of the HCV IRES agreed with its known structure^51,52^ when constrained by in-cell and *in vitro* DMS probing (Fig. 2a,b). The known structure reconstructed from Kieft et al. 1999^53^ diffiers at 4 nucleotides from the probed structure (annotated in Supp. Fig. 2a). The probed variant is fully functional in bicistronic and monocistronic luciferase reporters^20,54,55^. Discrepancies between the *in vitro* and in-cell DMS probing of HCV were identified by ΔDMS (see Methods). Most diffierences in-cell vs. *in vitro* did not disagree on base-pairing state: more reactive bases in cells were already modeled as unpaired. However, higher reactivities in cells were clearly visible around HCV domain IV, which contains its translation start codon (Fig. 2c).

**Figure 2.**
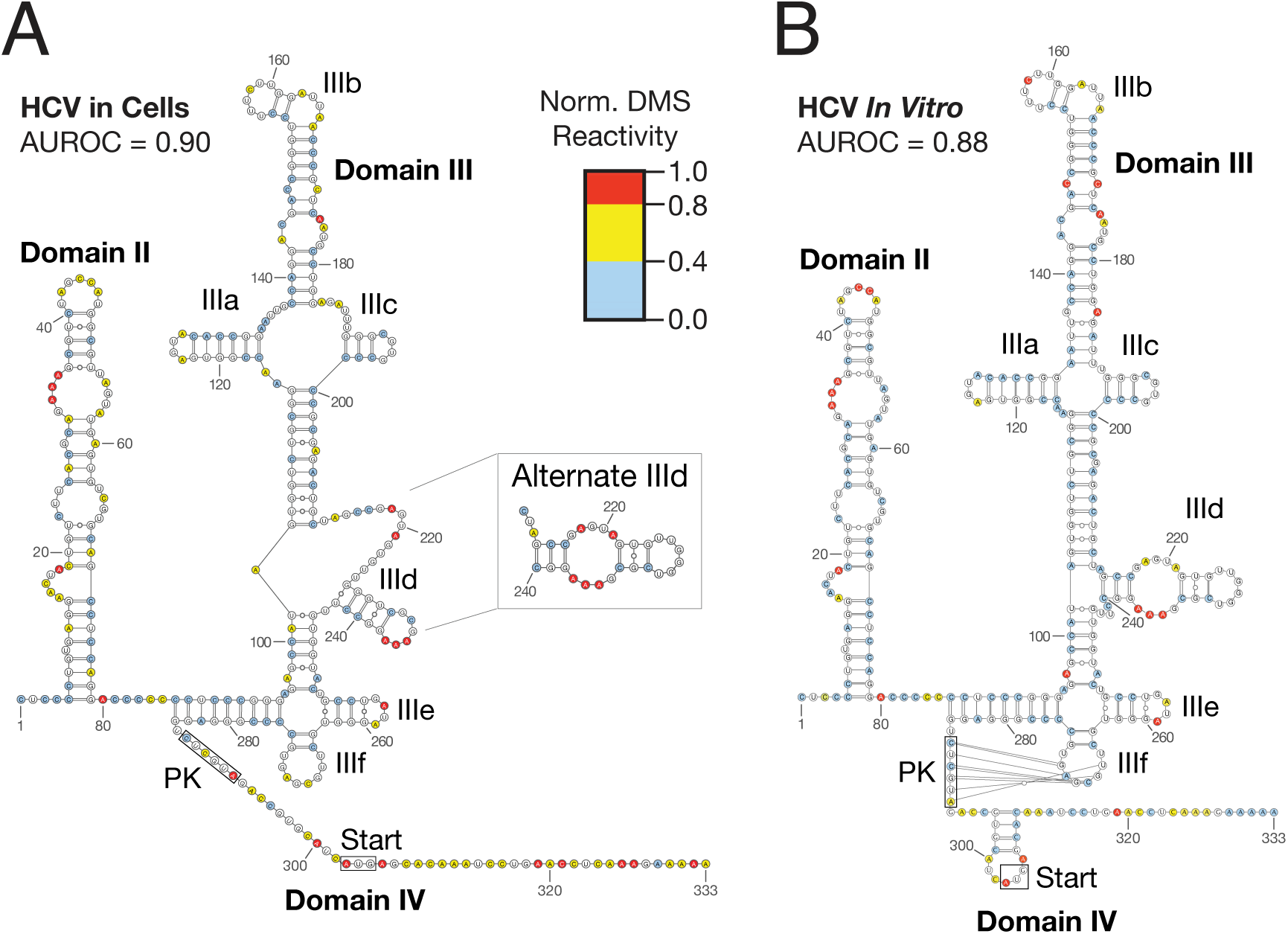
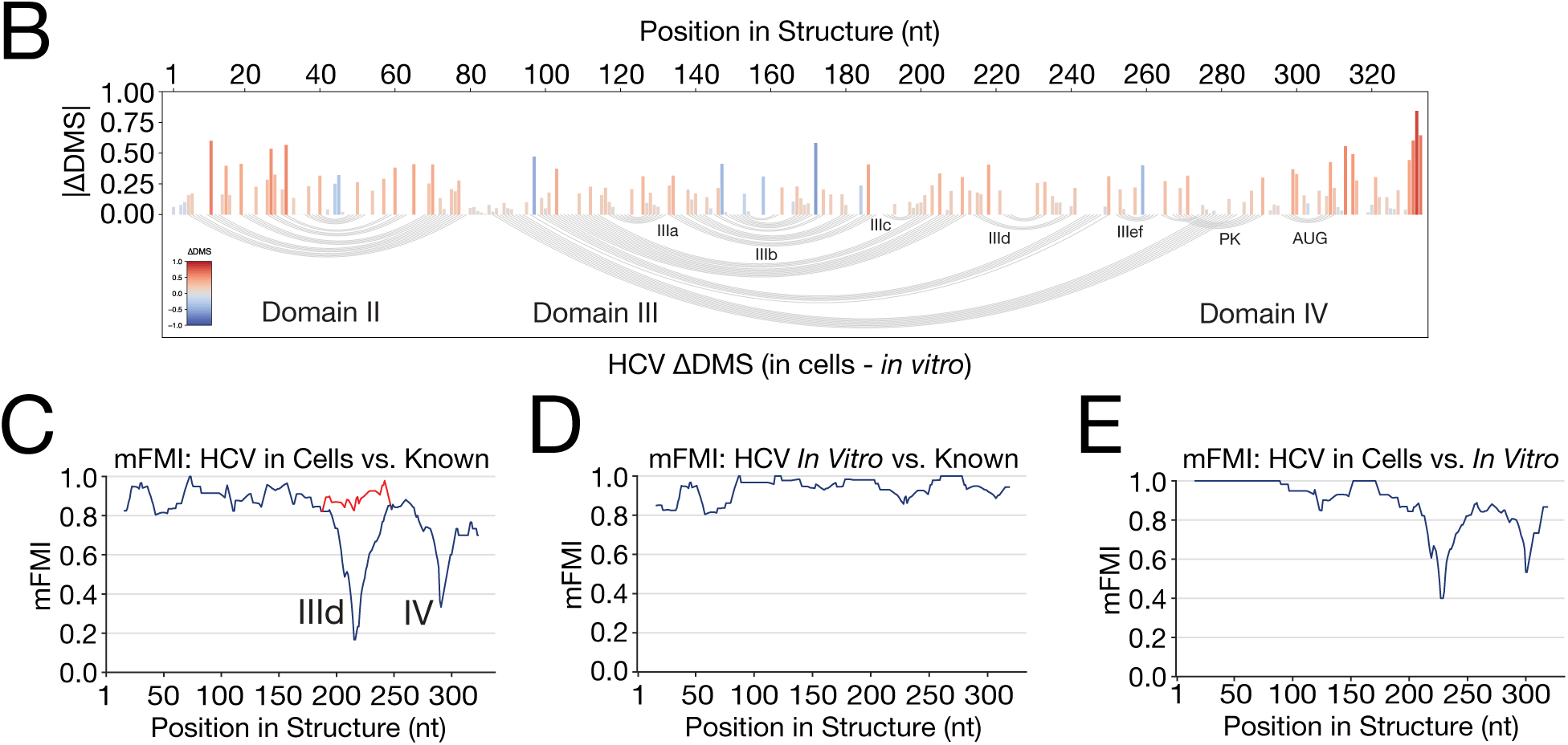
Hepatitis C virus (HCV) IRES RNA secondary structure with DMS-MaPseq. (**A**) The HCV 5’UTR IRES secondary structure constrained by DMS-MaPseq probing in HEK293T cells (AUROC = 0.90). Normalized DMS reactivities calculated as described. White: no data (Gs and Us). Blue: shielded bases, 0.0-0.4. Yellow: bases of intermediate reactivity, 0.4-0.8. Red: highly reactive bases, 0.8-1.0. Normalized reactivity scale and structure prediction parameters are identical in all figures (see Methods). Domains of HCV are labeled consistent with previous reports. An alternative prediction of domain IIId which more closely matches its known structure is inlaid. **(B)** The HCV IRES modeled from *in vitro* probing (AUROC = 0.88). **(C) Δ**DMS between the normalized in cell and *in vitro* probing data, with the model from Fig. 2b expressed as a corresponding arc plot (see Methods). ΔDMS is expressed as its absolute value and color coded on a scale of -1 to 1. **(D)** Modified Fowlkes-Mallows Index (mFMI, see Methods) quantitative structure comparison between the in-cell model and the known structure in Supp. Fig. 2a. Notable areas of disagreement in base-pairing labeled for domain IIId and domain IV. The red line corresponds to the mFMI of the inlaid alternative prediction. **(E)** mFMI between the *in vitro* model and the known structure. **(F)** mFMI between the in cell and *in vitro* models in Fig. 2a.

Agreement between our DMS-MaPseq constrained predictions and the known structure was assessed by modified Fowlkes-Mallows Index^37,40^ (mFMI, see Methods). mFMI on the in-cell structure indicated high agreement with the known structure of the HCV IRES except for domain IIId and domain IV. The inlaid alternative prediction of HCV domain IIId in Fig. 2a improves its agreement with the known structure, with the alternative mFMI shown in red (Fig. 2d). While this prediction more accurately captured domain IIId, it disrupted modeling of the 4-way IIIef junction, and given its high reactivity, DMS-constrained SHAPE-KNOTS^56^ did not identify the HCV pseudoknot (not shown). Domain IV appeared linear in our in-cell prediction (Fig. 2a). Unlike the in-cell model, *in vitro* constrained folding of HCV ubiquitously recapitulated its known secondary structure (Fig. 2b,e). mFMI comparing the in-cell and *in vitro* structures showed higher agreement in domain II than the comparisons to the known structure (Fig. 2f). Domain II harbors three of the four nucleotides that diffier from the probed structure (labeled in Supp. Fig. 2a).

Correspondence of DMS-MaPseq constraint data to the predicted structures was assessed by observing the area under the receiver operator characteristic curve (AUROC, see Methods). While AUROC does not measure RNA structure prediction accuracy, reliable predictions typically have AUROC > 0.80, indicating high agreement between signal and structure^37^. The HCV IRES prediction in cells has AUROC = 0.90 (Fig. 2a). DMS-MaPseq probing on the HCV IRES in cells was highly reproducible (n = 3), with R^2^ of 0.99, considering reactivity on As and Cs only (Supp. Fig. 1b, see Methods). The *in vitro* data on HCV (n = 2) was similarly reproducible with R^2^ = 0.96 (Supp. Fig. 1c). We compared the DMS-MaPseq data to the known structure. With in-cell DMS probing, AUROC = 0.76, indicating some disagreement, likely stemming from domain IV. *In vitro* DMS-MaP seq data agreed better with the known HCV structure with AUROC = 0.80 (Supp. Fig. 2a).

Identical folding parameters (see Methods) captured the known RNA secondary structure of the EMCV IRES, and the resulting model is consistent with previous reports based on *in vitro* probing and evolutionary conservation^57,58,59^. The EMCV prediction exhibited strong agreement with the DMS probing data with AUROC = 0.88 in-cell, and AUROC = 0.88 *in vitro* (Fig. 3a,b). Comparing *in vitro* DMS-MaPseq probing to in-cell, we observed a similar pattern in EMCV and HCV: ΔDMS (in-cell – *in vitro*) shows elevated normalized reactivities in cells surrounding the translation start codon in both viral IRESes (Fig. 3c). Outside this, neither appears to exhibit a substantially remodeled base-pairing state in the cellular environment compared to *in vitro* (Fig. 2c, Fig. 3c). Secondary structure modeling of the EMCV IRES in the cellular environment most closely resembled an *in vitro* model from Pilipenko et al. 1989^60^ which agreed best with a recent functional study^61^. This known structure spans from domain G to domain L, whereas the IRES probed in this report includes domains D-M. Comparing DMS signal-structure agreement on shared sequence with the known structure results in AUROC = 0.80 in cells and AUROC = 0.81 *in vitro* (Supp. Fig. 2b). The two models’ high similarity to the predicted structures is reflected by mFMI, with values > 0.8 across the UTR (Fig. 3d,e). DMS-MaPseq on EMCV was also highly reproducible, with R^2^ = 0.99 in cells (n = 2), and R^2^ = 0.87 *in vitro* (n = 2) (Supp. Fig. 1f,g). Overall, DMS-MaPseq is effective and reproducible for modeling large RNA secondary structures in cells and *in vitro*.

**Figure 3.**
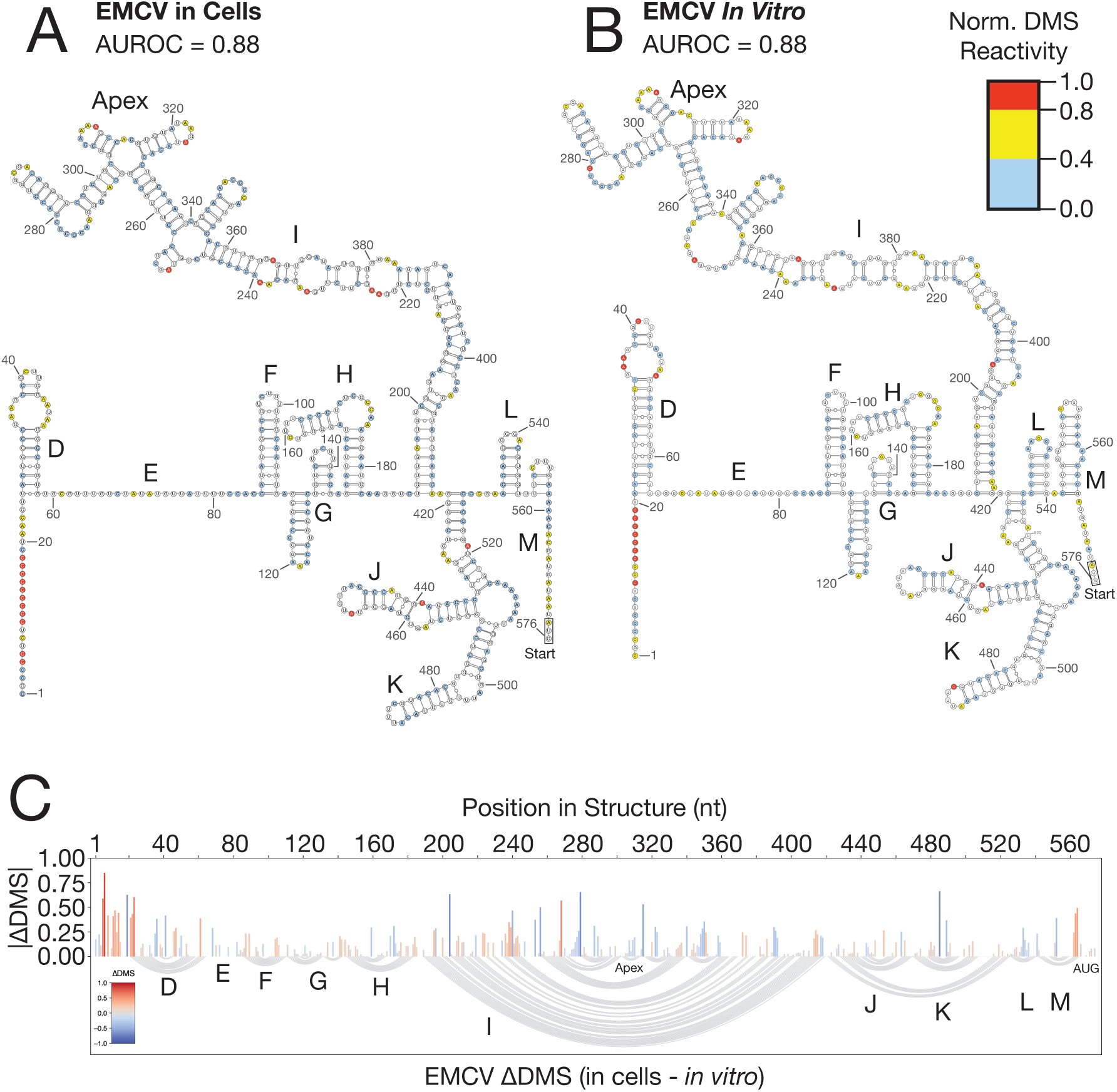
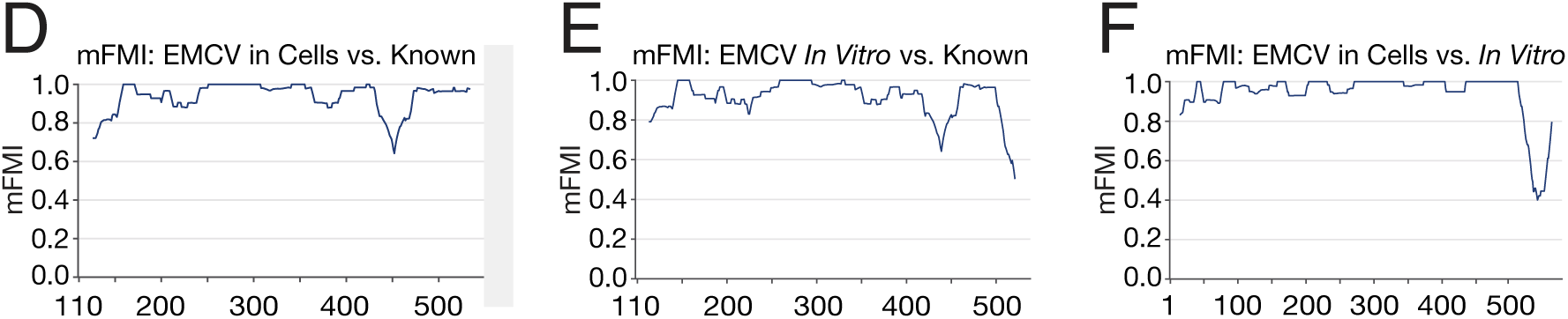
Encephalomyocarditis virus (EMCV) IRES by DMS-MaPseq. **(A)** The EMCV IRES secondary structure constrained by DMS-MaPseq, probed in HEK293T cells AUROC = 0.88). **(B)** Secondary structure constrained by *in vitro* probing of the EMCV IRES. (C) ΔDMS between in cell probing and *in vitro* probing of EMCV expressed as described, with the *in vitro* model in Fig. 3b depicted by an arc plot. **(D)** mFMI comparing in-cell EMCV (Fig. 3a) and the known structure in Supp. Fig. 2b across their shared sequence range. **(E)** mFMI comparing *in vitro* EMCV (Fig. 3b) with the known structure. **(F)** mFMI between in cell and *in vitro* models (Fig. 3a vs. Fig 3b).

### Modeling the RNA secondary structure of Mouse Insulin Receptor (Insr) 5’UTR constrained by DMS-MaPseq

To identify specific structural requirements for IRES activity, we modeled the RNA secondary structure of the Insr 5’UTR in cells and *in vitro*. DMS-MaPseq libraries were prepared from cultured cells and from *in vitro* reactions as described (Fig. 1a). These constraint data were applied with identical folding parameters to those used for the concurrently probed HCV and EMCV IRESes. The in-cell model had apparent regions of low base-pairing and agreed with the probing data with AUROC of Insr in cells = 0.87 (Fig. 4a). *In vitro*, a very similar structure was predicted, but all domains appear more paired than in cells (AUROC *in vitro* = 0.81, Fig. 4b). A notable difference is a register shift at 226-280 pairing with 290-340 *in vitro* instead of 246-280 pairing with 290-340 by the in-cell model. Comparing signal between in-cell and *in vitro* probing data by ΔDMS appears to indicate melting of the Insr 5’UTR at these positions in cells. Proximal to the AUG, ΔDMS also revealed a different pattern to the viral IRESes: normalized Insr DMS reactivity exhibited little change within 15nt of its translation start codon in cells compared to *in vitro* (Fig. 4c).

**Figure 4.**
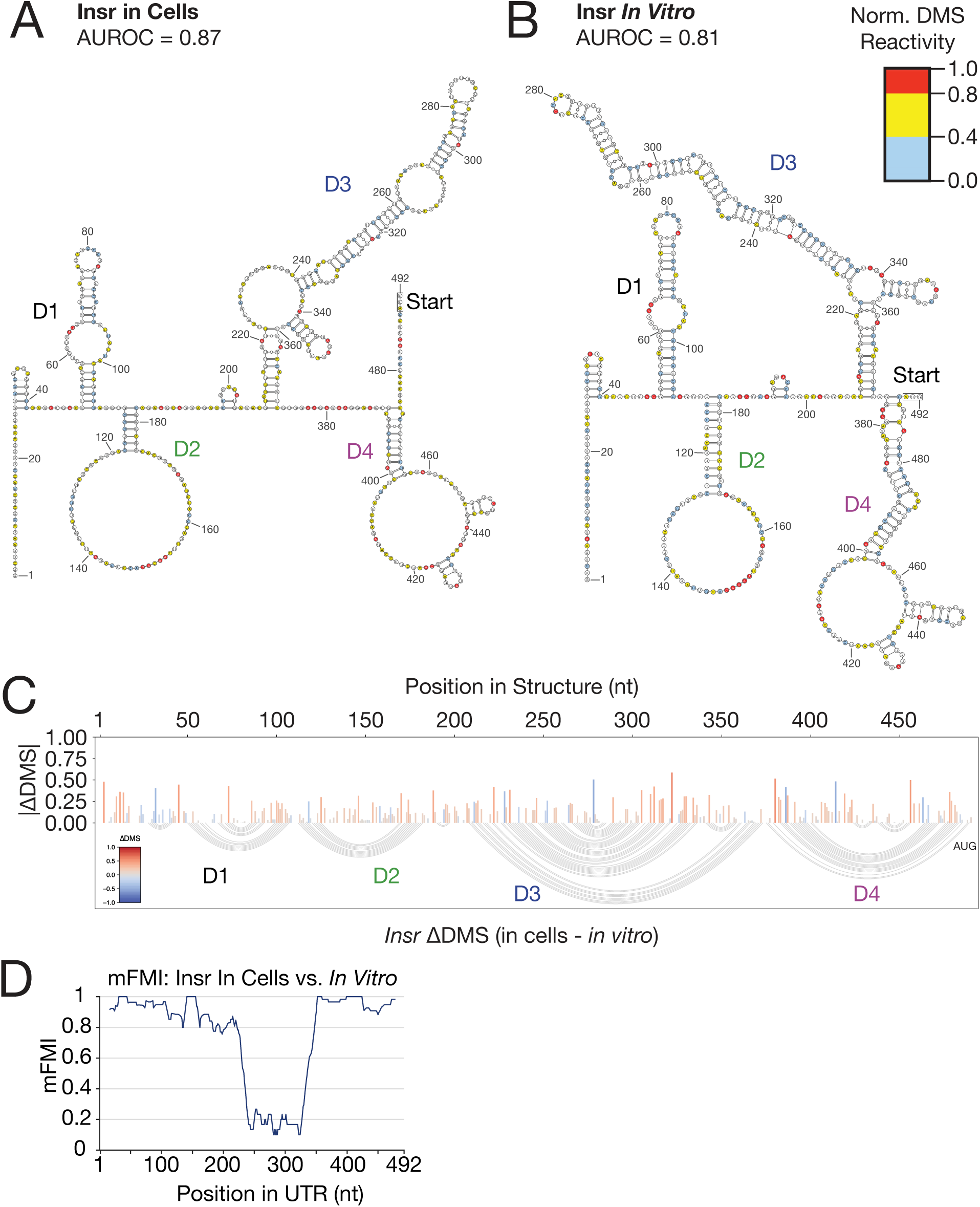
Modeling the RNA structure of the mouse Insr 5’UTR with DMS-MaPseq. **(A)** Model of the mouse Insr 5’UTR constrained by DMS-MaPseq in HEK293T cells. Exact domains divisions are outlined in Supp. Fig. 3a. **(B)** Model constrained by *in vitro* DMS probing. **(C) Δ**DMS comparison between in cell and *in vitro* probing. **(D)** mFMI between the predicted structures in A and B.

For ease of discussion, we assigned names: domain 1 (D1, nucleotides G1-U111), domain 2 (D2, G112-A207), domain 3 (D3, U225-C336), and domain 4 (D4, A382-U483). A diagram of these domains mapped to the *in vitro* model in can be found in Supplementary Figure 3a. Domain 1 is partially structured and is highly conserved. Domain 2 is largely modeled as unpaired. Domain 3 contains a stem loop with bulges, which exhibits a register shift *in vitro* vs. in-cells. Assessed by mFMI, the differences in structure between the two models centers on the register shift, with the rest of the structure maintaining mFMI > 0.8 (Fig. 4d). Domain 4 is most proximal to the AUG start codon with two predicted stem loops in its apex. There is little conservation in the mouse Insr 5’UTR outside of domain 1 (Supp. Fig. 3b). Following the area of high conservation, which extends to C138, there is a large stretch of unpaired A and C nucleotides that exhibit high mutation frequencies (C140-C208 in Fig. 4a). Probing of Insr was also highly replicable in cells with R^2^ = 0.98 and *in vitro* with R^2^ = 0.98 (Supp. Fig. 1j,k). We used these models to design a mutagenesis strategy aimed at identifying structural requirements for IRES activity within the Insr 5’UTR.

### Large deletions identify a region distal to the translation start AUG as critical for IRES-mediated translation initiation

To pinpoint the functionally relevant areas of the predicted structures for the Insr IRES, we began by introducing deletions based on the domains defined in our model of the structure. An initial assessment of the effects of these deletions on Insr IRES function was conducted using a bicistronic luciferase reporter system transfected into HEK293T cells. We have previously used this reporter system to investigate the relationship between the Insr IRES and several eukaryotic translation initiation factors^20,54,55^. The bicistronic system was chosen because monocistronic reporters of cellular UTRs may promote translation by both cap-dependent as well as cap-independent mechanisms *in vivo*. The bicistronic luciferase reporters have a cap-dependent Renilla luciferase ORF which is followed by the IRES in an intercistronic UTR that allows for translation of the downstream firefly luciferase ORF. This first cistron ensures the firefly ORF reports on internal ribosome entry, and it offers a built-in control for potential differences in RNA abundance arising from transfection efficiency or RNA stability. In our construct, the bicistronic transcript is driven by a relatively weak Rous Sarcoma Virus long terminal repeat (RSV LTR) promoter to better represent the low abundance of mouse Insr mRNA. A synthetic polyadenylation signal, as well as the RSV LTR polyadenylation signal, are included to limit transcripts originating elsewhere in the plasmid from extending into the bicistronic reporter.

In bicistronic reporters, the metric for IRES-mediated translation is firefly luciferase (FF) signal divided by Renilla luciferase (Ren) signal. The relative IRES capability of each mutant is expressed using their FF:Ren ratio as a percentage of that of full-length Insr UTR (IRES activity, see Methods). Mean IRES activity metrics across technical replicates from each experiment are plotted as data points in all charts. Each data point represents the value from a single biological replicate. Biological replicates were independently transfected on different days. Error bars are the standard deviation of the biological replicates, except for full-length, which is the standard deviation of technical replicates (see Methods). We utilized a non-IRES control containing beta-globin 5’UTR (BG), which has 3% or less IRES activity, as well as a positive control viral IRES (HCV), typically presenting at 40-60% IRES activity in our bicistronic reporters (Fig. 5a). BG provides a baseline to assess the performance of the assayed intercistronic UTRs in comparison to a cap-dependent UTR. HCV ensures translation does not generally differ across our assays. All plasmids generated for this study can be found in Table 2, and all raw luciferase assay data can be found in Table 3.

**Figure 5.**
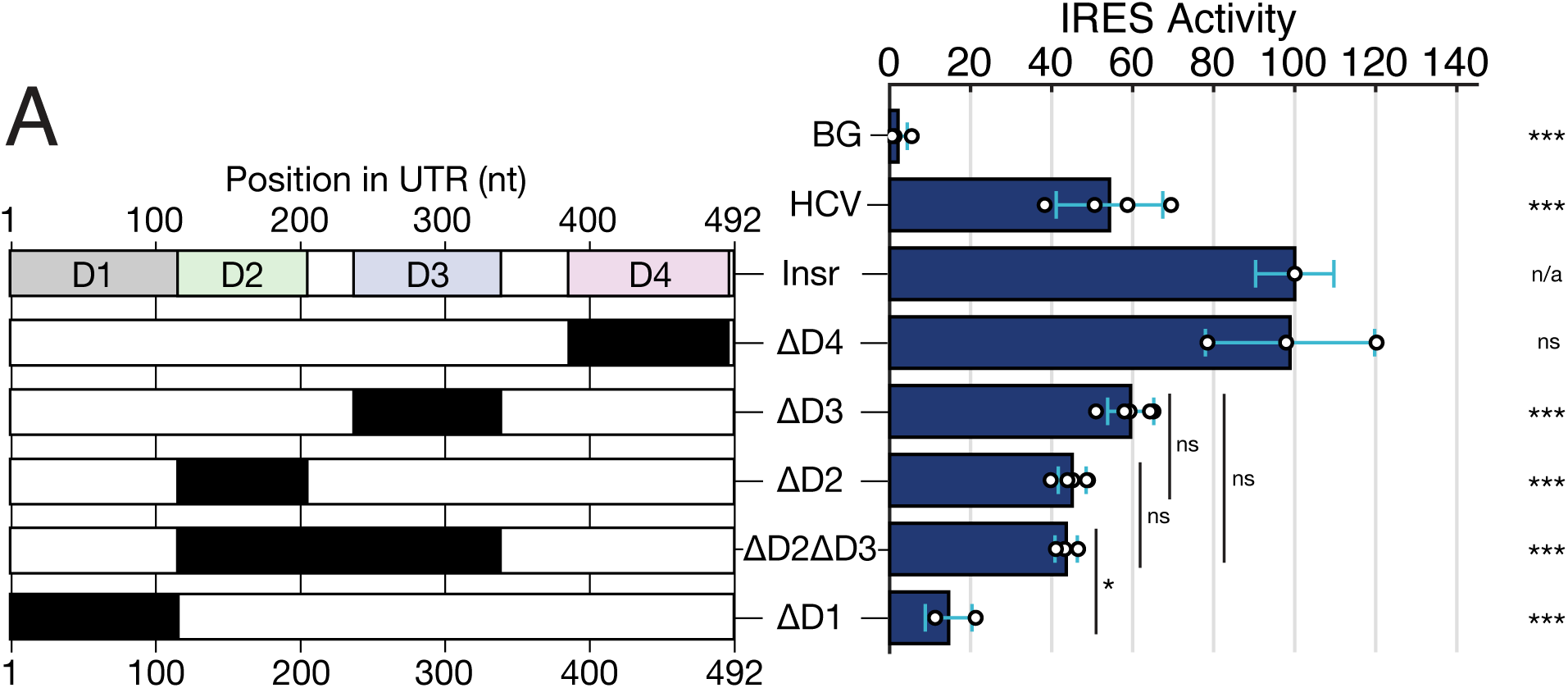
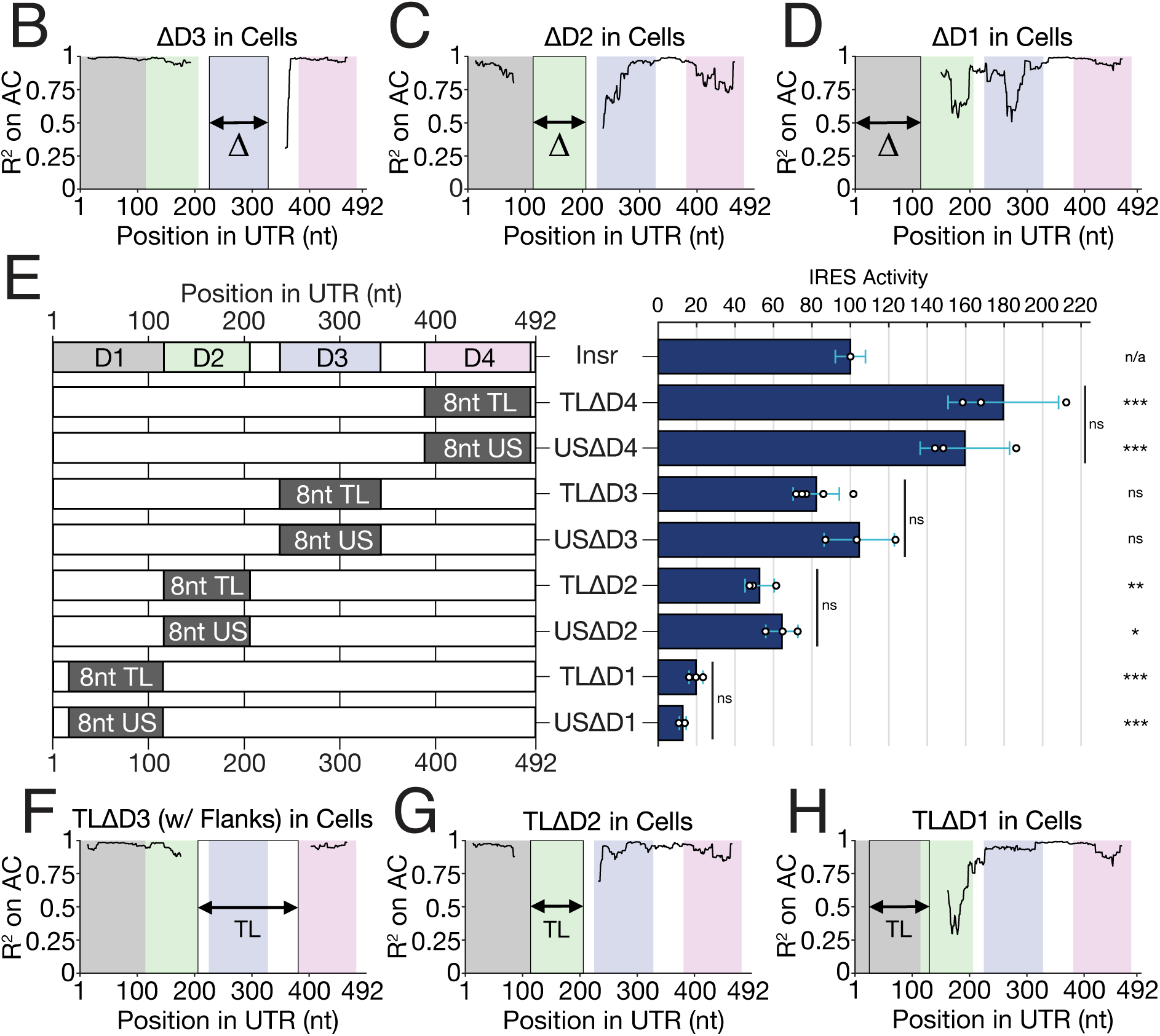
Identifying critical portions of the Insr 5’UTR. **(A)** Performance of bicistronic luciferase reporters deleting each indicated region. At left, black bars indicate deletions and white bars indicate that the sequence is retained. IRES activity is firefly/Renilla luciferase signal ratios divided by that of full length Insr 5’UTR (Insr). BG: cap-dependent UTR serving as negative control. HCV: viral IRES serving as positive control. Points: mean for each biological replicate, with three technical replicates each. Error bars: standard deviation of these means. Asterisks indicate p-values by ANOVA followed by Tukey’s HSD: * = p ≤ 0.05, ** = p ≤ 0.005, *** = p ≤ 0.0005. Asterisks at right are in comparison to full length Insr UTR. See Methods for complete details. **(B-D)** Coefficient of determination (R^2^) considering only signal on As and Cs, comparing independently probed mutants and the full length Insr 5’UTR (see Methods). Mutations are indicated. All probed in cells. **(E)** Bicistronic luciferase reporter performance of constructs with equivalent deletions to those in Fig. 5a, but with tetraloop hairpin (TL) or unstructured (US) replacements. **(F-H)** DMS signal on the tetraloop hairpin constructs compared to the full length Insr 5’UTR probed in cells. Regions discussed are color coded: D1, gray. D2, green. D3, blue. D4, magenta.

The deletions had varied effects on IRES activity. We assessed the effect on our reporters using analysis of variance (ANOVA) and Tukey’s HSD post-hoc test (see Methods). Surprisingly, a full deletion of domain 4 (ΔD4), which is most proximal to the translation start codon, has no effect (99±21%). Deletions of domain 3 (ΔD3, 60±6%) and domain 2 (ΔD2, 45±3%) damaged IRES activity, and were insignificantly different from one another. Combining the deletions of D2 and D3 (ΔD2ΔD3, 44±3%) did not reduce activity further. ΔD1 (15±6%) had the strongest effect observed among the large- scale deletions (Fig. 5a).

To assess whether the functional damage from deletions (ΔD3, ΔD2, and ΔD1) was a consequence of RNA secondary structure disruptions, we conducted DMS- MaPseq on mutants in cells and *in vitro*. Sequence deletions bring previously distal portions of RNA closer together, possibly changing surrounding RNA secondary structure and indirectly affecting IRES activity. We opted for a structure-naïve approach to assess changes by observing the signal correspondence on the adenosines and cytosines probed by DMS. This allows us to spot differences in base-pairing state regardless of the accuracy of the secondary structure model. We plotted the coefficient of determination (R^2^) calculated in sliding windows comparing only signal on As and Cs (see Methods). For comparisons, R^2^ < 0.8 was considered likely to diverge in pairing state. In cells, where a narrower range of structure is permitted^26^, deletions had a minor influence on DMS signal: ΔD3 did not appear to disrupt D1 or D4, but immediately adjacent residues did appear impacted (Fig 5b). Surprisingly, ΔD2 disrupted adjacent signal in D1 and D3, as well as D4 (Fig. 5c). ΔD1 disrupts portions of D2 and D3, but not D4 (Fig. 5d), which could contribute to the potent effect of ΔD1 by impairing proper folding. *In vitro*, deletions had a more accentuated impact on DMS signal. ΔD3 *in vitro* appeared similar to in-cells, ΔD2 appeared to more greatly disrupt D1, and ΔD1 disrupted structure across the UTR (Supp. Fig. 5a,c,e). These structural disruptions could present a confounding variable in determining the requirements for IRES activity if the impairment of IRES activity is exacerbated by changed pairing states outside the deletion of interest.

### Replacements rescue IRES activity in a subset of mutants

We sought to preserve the potential structural influence of each domain on the rest of the UTR. To this end, reporters were created which replace each deletion with tetraloop hairpins (TLs). Tetraloops are strong 4-base loops that assist folding of critical structure and tertiary interactions in rRNA and other large functional RNAs^62^. Replacing predicted stems with TL hairpins dispenses with the specific structure and sequence of the stem, substituting two basepairs topped with the tetraloop. The chosen TL was 5’– CCGAAAGG–3’, a GNRA tetraloop used in other studies of RNA structure^63,64,65^ with two CG basepairs as the stem. If the deletions functionally disrupt IRES activity by perturbing secondary structure, TLs could repair adjacent pairing and rescue the IRES activity in comparison to complete stem deletions. Each predicted domain was progressively replaced with TL hairpins. Diagrams of the positions of these TLs on both Insr 5’UTR models can be found in Supplementary Figure 4b and 4c. The effects of these mutations were assayed by bicistronic luciferase reporters as previously described.

A TL hairpin in place of D1 (TLΔD1, 20±4%) functionally matched the effect of ΔD1, and TL replacement of the low pairing ΔD2 (TLΔD2, 53±8%) was not significantly different to the effect of ΔD2 (Fig 5a,e). However, domain 3’s TL (TLΔD3, 82±12%) exhibited a significant functional rescue compared to ΔD3 while removing the same sequence range. Surprisingly, a complete TL hairpin replacement of domain 4 (TLΔD4, 180±29%) significantly enhanced the activity of the UTR, but was far more variable than any other mutant tested (Fig. 5e). Only TL hairpins which reached the base of the predicted stems exhibited the effects depicted in Fig. 5e. Notably, other TL hairpins replacing part of D1 showed no significant impairment of activity (D1-TL1, 66±4%, D1- TL2, 81±13%, and D1-TL3, 80±9%). D1-TL3 precisely replaces the large stem loop at G50-C108. Similarly, the effect of TLΔD2 was not observed in partial replacements: mimicking the predicted helix with a TL (G112-C118 pairing with G176-C182) had no significant effect (D2TL-3, 74±9%), and shifted TL replacements which impact one half of this helix also had no effect (D2TL-1, 106±22%, and D2TL-2 74±27%). In D3, TL hairpins rescued compared to ΔD3 regardless of their exact position (D3-TL1, 101±14%, D3-TL2, 94±25%, and D3-TL3, 91±13%). Finally, incomplete replacement of D4 (D4- TL1, 102±19%) had no effect (Supp. Fig. 4a).

We surveyed the TL hairpins’ influence on RNA secondary structure using DMS-MaPseq. In cells, TLΔD3 was very similar to ΔD3, with intact DMS signal across the UTR (Fig. 5f). TLΔD2 slightly enhanced agreement with the full-length structure compared to ΔD2 (Fig. 5g). TLΔD1 did not fix the structural disruption caused by ΔD1 in D2, but did appear to rescue its influence on D3 (Fig. 5h). *In vitro* probing however, showed significant disruptions in both adjacent and distal domains (Supp. Fig. 5b,d). Despite the tetraloop hairpin in place of D3 offering a functional rescue in bicistronic luciferase reporters, TLΔD3 disrupts the border between D1 and D2 in *in vitro* (Supp. Fig. 5b). Similarly, TLΔD2 further disrupted adjacent UTR *in vitro* compared to ΔD2 (Supp. Fig. 5d). Overall, reactivity across the UTR remained largely intact in cells, with some assistance from TL hairpins. Significant structural disruptions were observed *in vitro* that increased with the addition of TL hairpins, despite the functional enhancement of TLΔD3 and TLΔD4 compared to equivalent broad deletions.

We created linear linker mutants to investigate whether the TL hairpins were structurally contributing to the function of the Insr 5’UTR. We replaced TL hairpins with CAA repeats of equal length (8nt, 5’–AACAACAA–3’). These linkers are reliably linear in RNA structures^66^ and are denoted in this report by the ‘unstructured’ prefix rather than TL (USΔD). If rescue of function by TL hairpins contributes structural support to the UTR, their unstructured counterparts will not rescue. The US mutants showed an identical capacity for IRES activity as their TL hairpin equivalents, with USΔD4 (160±23%) enhancing activity, USΔD3 (104±18%) rescuing to be indistinguishable from full-length UTR, USΔD2 (64±8%) offering no rescue, and USΔD1 (13±2%) causing equivalent damage to TLΔD1 and ΔD1 (Fig. 5e). Additionally, disruptions in predicted helices in each domain did not affect the reporter (Supp. Fig. 6a). The effects observed in the tetraloop mutants do not depend on the tetraloops’ structure but do rely on added nucleotides in place of the predicted stems.

### Constructing a minimal sufficient mutant of the Insr 5’UTR IRES

Finding the minimal components for IRES activity would allow us to more precisely investigate its function by providing a less complex sequence element than the full-length UTR. Informed by the importance of domain 1, we created a series of sufficiency constructs to assess whether a structurally intact D1 is sufficient for IRES. First, each of the domains were individually assayed for sufficiency using the constructs D4-Solo (1±1%), D3-Solo (1±1%), D2-Solo (3±1%), and D1-Solo (10±2%). None exceeded 5% the full-length UTR’s IRES activity except D1-Solo (Fig. 6a and Supp Fig. 3b). We suspected that impairment of D1-Solo may be due to compromised structure. However, DMS-MaPseq in cells on D1-Solo exhibited only a minor decline in signal correspondence, and much of D1 appeared intact with R^2^ > 0.8 (Fig. 6b). We then added back portions of the UTR to achieve sufficiency, starting with only the conserved area (up to C138). The conserved sequence alone (Cons-Solo, 28±1%), improved activity. Adding back the rest of D2 (D1D2, 49±14%) further increased activity (Fig. 6a). By DMS-MaPseq, D1D2 appeared to recapitulate the signal on both D1 and D2 (Fig. 6c). While D1D2 is critical, it is not fully sufficient for full IRES activity, even when structurally intact. We suspected adding back a TL hairpin in place of D3 would rescue the activity of D1D2 like TLΔD3 rescued in the full-length context (Fig. 5a,e). As predicted, in D1D2TL (78±11%), a TL hairpin in place of D3 rescued activity compared to D1D2 alone (Fig. 6a). However, D1D2TL is still significantly impaired compared to full-length UTR. (28±6%). This set of mutants indicate a requirement for D1 and D2, and a graded sufficiency as generic elements are added back in place of D3 and D4.

**Figure 6.**
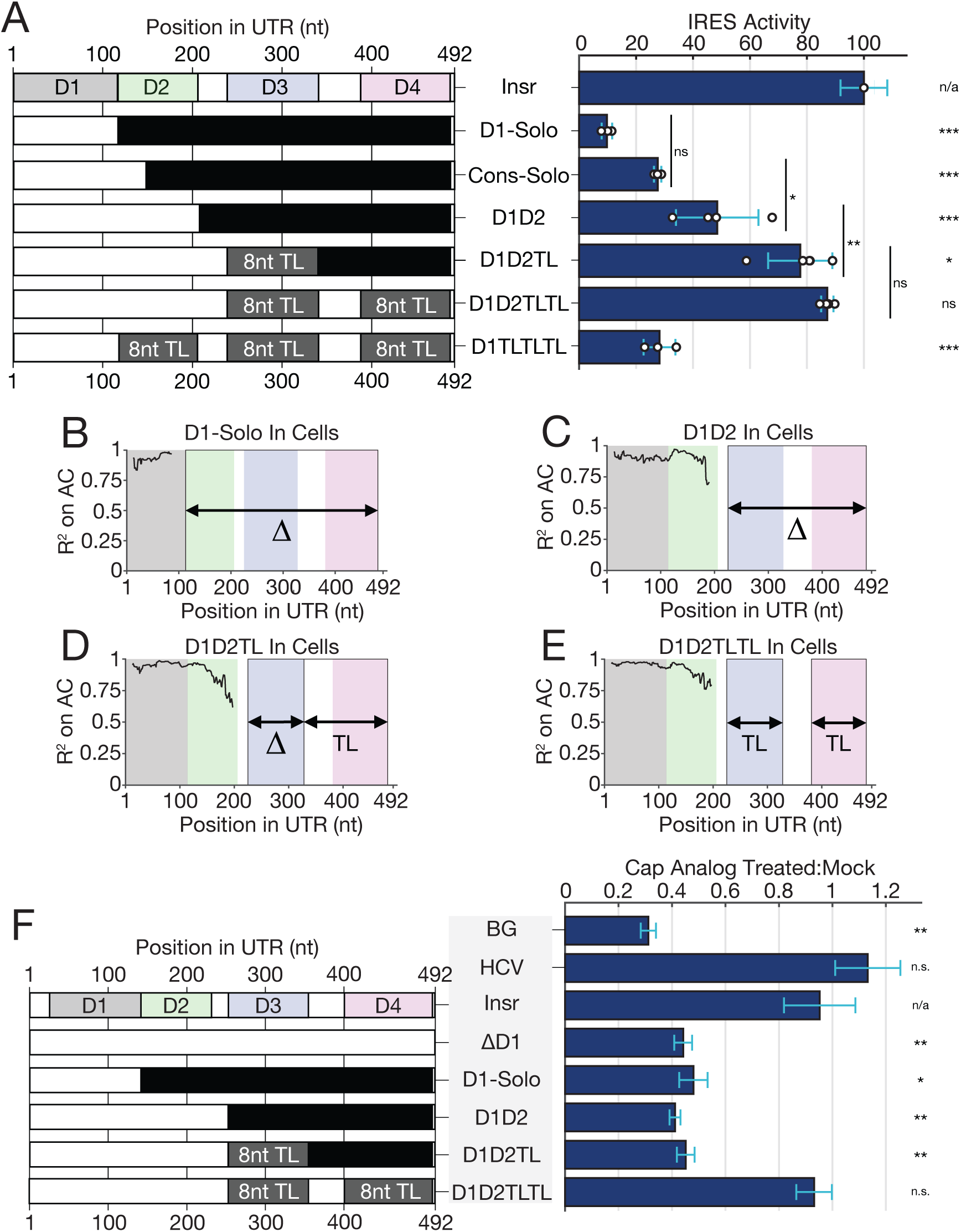
The minimal sufficient mouse Insr 5’UTR IRES. **(A)** Bicistronic luciferase reporters’ performance on sufficiency mutants with metrics and statistics as previously described. **(B)** Comparing DMS-MaPseq on these sufficiency mutants with R^2^ plots produced as previously described. Deletions and TL hairpin reactions. For this experiment, n = 2. Error bars: average standard deviation from technical replicates.

Given this result, we anticipated adding a TL hairpin in place of D4 to D1D2TL would also behave the same as the full-length context, offering an enhancement to activity (Fig. 5e). Surprisingly this mutant, D1D2TLTL (87±2%), did not significantly enhance activity compared to D1D2TL, but did exhibit full sufficiency (Fig. 6a). In-cell DMS-MaPseq was conducted on the D1D2TL and D1D2TLTL, and both had a strong overall correspondence to full length UTR (Fig. 6d,e). When probed *in vitro*, D1D2TLTL appeared to have a similar pattern of disruptions to TLΔD3 (Supp. Fig. 5f). In the context of D1D2TLTL, replacement of D2 with a TL hairpin (D1TLTLTL) greatly impaired the IRES (28±6%). This set of mutants indicate a requirement for D1 and D2, and a graded sufficiency as generic elements are added back in place of D3 and D4.

### Monocistronic luciferase reporters challenged with cap analog in vitro corroborate the minimal sufficient Insr 5’UTR IRES

To corroborate our minimal sufficient 5’UTR identified by bicistronic reporters, we generated monocistronic mRNAs *in vitro* and assayed their activity in rabbit reticulocyte lysate (RRL). *In vitro* translation allows for direct treatments against cap-dependent translation initiation. Challenging the translation extract with excess N^7^- methylguanosine (m^7^G) competes with mRNA cap binding of eukaryotic translation initiation factor 4E (eIF4E), inhibiting its ability to recruit transcripts to ribosomes^67,68^. Transcripts dependent on their 5’ caps to initiate translation are susceptible to this treatment^69^. Monocistronic firefly luciferase reporter mRNAs were translated *in vitro,* and signal was measured after 30 minutes of incubation in RRL. The response of the constructs to m^7^G cap treatment was assessed in the context of untreated RRL, where other mRNAs were present to assay their performance under competitive conditions. For this experiment, n = 2. Two controls were used to assess nonspecific effects of m^7^G on the extract: the BG 5’UTR reporter measures the overall effect on cap-dependent translation initiation and benchmarks the performance of transcripts with no IRES activity. The HCV IRES reporter controls for non-specific effects of cap analog treatment on the translation of cap-independent transcripts.

Addition of 250µM m^7^G cap analog diminished translation of BG to 0.31±0.03 (Cap:Mock), whereas signal from the positive control HCV IRES and full length INSR were unaffected. The ΔD1 mutant (0.44±0.03) as well as D1-Solo (0.48±0.05) saw substantial loss of activity under cap treatment compared to mock treated comparable to the effect seen on BG. This recapitulates the results of the bicistronic reporters in cells where they were among the most impaired constructs for IRES activity. Interestingly, the graded sufficiency of D1D2 and D1D2TL seen in cells was not evident *in vitro*: instead, both D1D2 (0.41±0.02) and D1D2TL (0.45±0.03) were significantly impaired. D1D2 and D1D2TL were highly susceptible to m^7^G cap treatment, with Cap:Mock signal ratios insignificantly different from ΔD1 and D1-Solo. By contrast, the minimal sufficient D1D2TLTL (0.93±0.06), with tetraloop hairpins replacing domains 3 and 4, appeared entirely sufficient and insignificantly different from full length Insr 5’UTR *in vitro* (Fig. 6f), in agreement with its sufficiency in bicistronic reporters.

## DISCUSSION

Previous work from our lab demonstrated that insulin receptor 5’UTR allows cap-independent translation and likely contains an internal ribosome entry site (IRES). This capability is conserved from insects to mammals^19,20^. The RNA secondary structure of mouse Insr 5’UTR and this structure’s role in cap-independent translation initiation was unknown. We applied DMS-MaPseq to gain insight into the RNA structure of *Mus musculus* Insr 5’UTR in cells, and we validated our strategy by probing two viral IRESes of known structure (HCV and EMCV) alongside the Insr IRES in the same cells (Fig. 1). In-cell DMS-MaPseq data constrained secondary structure models of the HCV and EMCV viral IRESes which agree well with their known *in vitro* structures (Fig. 2, Fig. 3). For HCV, our result agrees with a recent report probing RNA secondary structure of the HCV viral IRES in the context of the viral genome in cells^31^. To our knowledge, this is the first report of the EMCV viral IRES structure in cells. Here we present a model of the secondary structure of the Insr 5’UTR informed by DMS chemical probing in cells and *in vitro* (Fig. 4) which we used to generate structure informed alterations in the RNA. These experiments identify a conserved portion of the Insr 5’UTR critical for its IRES activity (Fig. 5). This knowledge facilitated our design of a minimal sufficient IRES (Fig. 6).

Our results demonstrate the effectiveness of DMS-MaPseq for modeling RNA secondary structures larger than 300nt. Both in-cell and *in vitro* DMS probing allowed for accurate prediction of the HCV IRES secondary structure. The 333nt HCV IRES has three domains (II, III, IV) and features a 4-way junction (IIIe-f) which interacts with a 6bp pseudoknot critical for positioning the start codon AUG contained in domain IV^14,64^. In cells, default folding parameters captured most of the structure, but stem loop IIId was more accurately modeled in the second lowest energy prediction (Fig. 2a). *In vitro*, the entire secondary structure was modeled accurately (Fig. 2b). Our observation of a highly DMS-reactive pseudoknot and domain IV is specific to in-cell DMS probing. High ΔDMS signal adjacent to the AUG (Fig. 2c) has a consequent effect on folding of domain IV, quantified by mFMI (Fig. 2d-f). This indicates that this region is biologically linearized, likely by translation. Additionally, the HCV IRES attaches to 80S ribosomes translating other RNAs, which may affect its secondary structure; previous analysis of the HCV IRES bound to translating 80S ribosomes *in vitro* excludes domain IV^70^. The highly reactive sequence begins at C288 (AUG -15), which corresponds to previous reports using DMS toeprinting of the ribosome on the HCV IRES^71^ *in vitro*. Our DMS-MaPseq constraint data agree strongly with the known structure of the HCV IRES, but HCV is 159nt shorter than the 492nt mouse Insr 5’UTR. The difficulty of RNA secondary structure prediction scales with size, as greater base-pairing possibilities widen the range of *in silico* RNA structure prediction^72^.

To assess our strategy with a structure more analogous in size, we probed the 576nt EMCV IRES in cells for the first time, and we probed *in vitro* for comparison. The consensus structure for EMCV 5’UTR, shared across closely related viruses, contains fourteen stem loop domains, denoted A-N^57^. The EMCV IRES probed in this report contains domains D through M. Through repeated investigations, all but domain I have been resolved^60,59,61^. Chemical probing data does not perfectly agree with any particular model of domain I, but a recent functional study has refined some aspects of the active secondary structure^61^. In-cell modeling of the unresolved domain I most closely agrees with an earlier model of EMCV IRES^60^, which is preferred by the aforementioned functional study (Fig. 3a and Supp. Fig. 2b). *In vitro*, some structural differences were noticeable, with an extended stem loop at domain D, and a different pairing state below the apex of domain I (Fig. 3b). EMCV assessed by ΔDMS reflected the difference in pairing state seen in domain I. EMCV exhibited a similar pattern of ΔDMS to HCV: in-cell data had higher normalized reactivities than *in vitro* data surrounding the translation start codon (Fig. 3c). This likely means that a significant population of probed EMCV is translationally active. Unlike HCV, EMCV’s start codon is not contained in an RNA structure, though its relationship to structure does matter^73^. All depicted structures of EMCV exhibit high signal-to-structure correspondence by AUROC, denoted for each structure (Fig. 3a,b and Supp. Fig. 2b). Overall, EMCV IRES secondary structure prediction constrained by DMS-MaPseq agreed with its known structure both in cells and *in vitro* assessed by mFMI (Fig. 3d-f). We conclude that DMS-MaPseq is capable of accurately modeling large scale RNA secondary structures in cells and *in vitro*.

In the HCV^74,75,76^ and EMCV^60,77^ viral IRESes, conservation corresponds to the criticality of secondary structure as well as sequence-based interactions with translation machinery. This appears to be generally true of viral IRESes, but whether cellular IRESes have conserved structures is not yet clear^72^. The Insr 5’UTR exhibits little sequence conservation among mammals after the first 138nt (Supp. Fig. 3b). It is possible that this high sequence divergence among mammals reflects a low structural stringency for cap-independent translation initiation. We initially suspected that the high degree of transcript conservation on the 5’ end of the Insr 5’UTR was unrelated to IRES activity, as this region of the genome contains binding sites for factors that control Insr transcription^24^.

We sought to discern whether RNA secondary structure is responsible for Insr IRES activity by taking advantage of the strengths of DMS-MaPseq, which allowed us to start from a full-scale model of the Insr 5’UTR. The range of allowed RNA structures in cells is narrower than the energetically permissible secondary structures *in vitro* due to ATP-dependent processes such as translation and helicase activity^26^. *In vitro* probing allowed us to observe the intrinsic folding properties of the Insr 5’UTR absent the influence of RNA binding proteins, translation, and helicases. Modeling the Insr 5’UTR in cells and *in vitro* resulted in similar domain placement and architecture (Fig. 4a,b). Our comparisons by ΔDMS found that the Insr 5’UTR is generally less structurally intact in cells than *in vitro*. Unlike the viral IRESes, we did not observe elevated ΔDMS within 15nt of the translation start codon in Insr (Fig. 4c). Insr may be translated cap-dependently more frequently than the viral IRESes in our reporters, linearizing its UTR more often, which may contribute to the shifted register and ΔDMS profile of domain 3 reflected in mFMI between in-cell and *in vitro* (Fig. 4d). This suggests that the Insr IRES operates by a different mechanism than the probed viral IRESes. The Insr IRES may be less strictly reliant on specific RNA secondary structure, in agreement with its low evolutionary conservation. We sought to pinpoint which portion of the Insr 5’UTR is responsible for IRES activity by introducing structure-informed sequence changes to the RNA.

We first identified critical elements of the Insr IRES by screening the effects of mutations using bicistronic reporters in cultured cells. Broad domain deletions revealed a surprising area of criticality, distal from the translation start codon: removing domain 1 (ΔD1, 15±6%) had the greatest impact on the reporter (Fig. 5a). This location has also been implicated in the translation of human insulin receptor which contains a closely related sequence^78^. Our results indicate that this region may be conserved due to its effects on translation as well as its known impact on transcription^24^. In contrast to our deletions distal from the AUG, deletion of the domain most proximal to the AUG (ΔD4, 99±21%) had no effect, indicating that this IRES is unlikely to stringently load the translation start codon like the HCV viral IRES. Deleting the unpaired D2 (ΔD2, 45±3%) and the largely paired D3 (ΔD3, 60±6%) impacted the reporter similarly, and a combined deletion (ΔD2ΔD3, 44±3%) did not reduce activity further (Fig. 5a). The result from the combined deletion suggests D2 and D3 assist the IRES by a similar mechanism. This is surprising given D2 and D3’s substantially different RNA secondary structure: our predictions indicate that D3 contains the largest stem, but D2 is largely unpaired (Fig. 4a,b).

Due to their similar effect, we initially suspected that both ΔD2 and ΔD3 could affect proper folding of the critical D1, and we probed these mutants with DMS-MaPseq in cells and *in vitro*. This revealed differential disruptions in structure across the affected mutants that were more accentuated *in vitro* than in-cells (Fig. 5b-d, Supp. Fig. 5a,c,e). To rescue these disruptions, we introduced tetraloop hairpins to mimic the influence of each stem on adjacent RNA structure. Tetraloop hairpins at the apex of each domain had little effect on the reporter (Supp. Fig. 4a), suggesting that the functionally relevant elements of the UTR lie at the bases of the predicted stems. Tetraloop hairpins equivalent to the deletions positively affected the reporter only in domains 3 and 4. Domains 1 and 2 were not rescued by TLs or unstructured linkers, directly implicating them in the UTR’s function (Fig. 5e). Subsequent DMS-MaPseq on the TL hairpin replacements revealed some rescue in cells (Fig. 5f-h), but greater disruption *in vitro* (Supp. Fig. 5b,d,f), where the intrinsic folding properties of the UTR dominate. These structural disruptions *in vitro* did not correspond to the functional impact of TL replacements on our reporters. TL replacement of D3 (TLΔD3, 82±12%) rescued IRES mediated translation, but replacement of D2 (TLΔD2, 53±8%) did not (Fig. 5e). In-cell probing of the TL hairpin constructs revealed that the UTR’s allowed structure in cells is largely intact in both TLΔD3 and TLΔD2 (Fig. 5f,g). TLΔD1 rescues the disruption in D3, but not D2 (Fig. 5h), but exhibits similar functional impairment to ΔD1, supporting the conclusion that D1 is directly involved in the IRES.

After observing little correspondence between the structural and functional rescues of TL hairpins compared to broad deletions, we tested whether the structure of the TLs mattered for function by introducing linear linkers of equal length in their place. The TL hairpins’ structure was not relevant to the activity of the 5’UTR: linear linkers of equivalent length recapitulate their effects, including the rescue in place of D3 and the enhancement in place of D4 (Fig. 5e). This supports the idea that overall structure in the Insr 5’UTR is not responsible for its cap-independent translation capacity. This conclusion is also supported by the fact that disruptions in helices in each of the predicted domains have no effect (Supp. Fig. 6a). It is possible that a low degree of base-pairing facilitates interactions with IRES *trans*-acting factors as has been proposed for the human insulin receptor 5’UTR^78^. Alternatively, the Insr IRES may operate by scanning after internal entry like the poliovirus IRES, which scans a short distance and depends on upstream attachment elements^73^. This may require the unpaired elements adjacent to the stem loop in D1, which could be interrupted by tetraloop hairpin replacement. Alternatively, the position of predicted stem loops relative to one another, such as the proximity of D3 to D1, might interrupt IRES function. This could be generically rescued by linkers in place of a deletion, regardless of their structure.

Although D1 is necessary for complete IRES activity in the mouse Insr 5’UTR, D1 in isolation (D1-Solo) is not sufficient. D1-Solo is sufficient for structure in cells: it matches the signal on As and Cs seen in D1 in the full-length UTR (Fig. 6b). Because ΔD2 (45±3%) results in only a 2-fold effect (Fig. 5a), we expected D1 to be approximately 50% sufficient for Insr 5’UTR IRES function. While D1 is the most sufficient of any single region investigated, D1-Solo (10±2%) is still impaired 10-fold. The lack of sufficiency in D2-Solo (3±1%), D3-Solo (1±1%), and D4-Solo (1±1%) (comparable to beta-globin, also 1±1%) indicates that there are not independently sufficient sections of 5’UTR (Supp. Fig. 6b). Adding back UTR elements, starting with the conserved portion of D2, exhibits a graded improvement in sufficiency, with TL hairpin replacements of D3 and D4 fully sufficient (Fig. 6a). No structural property of these mutants explains their differences in function: D1-Solo (10±2%), D1D2 (49±14%), D1D2TL (78±11%), and D1D2TLTL (87±2%) all fold correctly in cells (Fig. 6b-e). In fact, D1D2TLTL, functionally indistinguishable from full-length UTR, appears significantly disrupted in part of D1 and in D2 *in vitro* (Supp. Fig. 5f). However, this disruption of intrinsic folding *in vitro* is not reflected by in-cell probing and is likely not allowed in the cellular environment (Fig. 6e). These results indicate that the structures and sequences of both D3 and D4 are dispensable, consistent with their lack of conservation, but that the context of D1 and D2 within the 5’UTR is key. The graded sufficiency of our sufficiency mutants may arise from distance between binding sites for translation machinery to the translation start codon, or from another influence on productive start codon selection. Whether the Insr 5’UTR IRES functions by direct placement of the AUG start codon (like the HCV IRES^14,64^) or by preinitiation complex scanning after the 40S ribosomal subunit binds (like the poliovirus IRES^79^) is unknown. The conserved portion of the Insr 5’UTR is sufficient for low level IRES activity, but it requires a contribution from the remainder of the UTR to achieve the activity of full length Insr 5’UTR.

We found that the minimal sufficient RNA, D1D2TLTL, behaves equivalently to the full-length UTR across translation contexts. We corroborated effects observed with our bicistronic reporter constructs with monocistronic luciferase reporters assayed using *in vitro* translation in untreated rabbit reticulocyte lysate (RRL). By adding an excess of m^7^G cap analog, the cap binding protein eIF4E is inhibited from recruiting transcripts to the ribosome^80^. Transcripts capable of cap-independent translation initiation are not impaired by this treatment, while cap-dependent transcripts have reduced translation capacity^81^. The cap-dependent BG control remained translated at 0.31±0.03 (Cap:Mock)

of the mock treated condition, which agrees with the established effects of cap analog treatment both on ribosome recruitment^67^ and the resulting effect on translation in RRL^82^. RRL is capable of translating uncapped RNAs that do not contain IRESes, albeit with substantially diminished translation activity^83^. As such, we considered a 70% reduction the maximum expected effect on compromised mutants. The ΔD1 RNA (0.44±0.03) and the D1-Solo RNA (0.48±0.05) were both highly susceptible to the cap analog treatment, indicating compromised IRES capability and recapitulating the necessary but insufficient nature of D1 *in vitro*. Surprisingly, there was no intermediate sufficiency of D1D2 (0.41±0.02) and D1D2TL (0.45±0.03) in this paradigm: both appeared entirely insufficient, and insignificantly different from ΔD1 and D1-Solo (Fig. 6f). This result contrasts the cell-based bicistronic assay. This could be a consequence of the different context of the monocistronic *in vitro* translation experiments from the bicistronic in cells experiments: the assayed UTR is in a different reporter context, operating on a different timescale, with a different organism’s translation machinery. While these differences may contribute to our results on reporters of intermediate sufficiency, they bolster the results on D1D2TLTL (0.93±0.06), which was also fully sufficient in bicistronic context in cultured cells. The conserved D1D2 is necessary and sufficient when distal from the translation start codon, and D1D2TLTL is sufficient across translation contexts.

## CONCLUSIONS

DMS-MaPseq revealed the RNA secondary structures of the HCV and EMCV viral IRESes in cells, which largely agree with their known structures *in vitro*. The site of translation initiation in both HCV and EMCV was linearized, visible by comparing DMS probing *in vitro* to in cells. Mutational profiling facilitated a new strategy for investigating the mouse Insr 5’UTR IRES: probing the full-scale structure and subsequently designing mutations to investigate its function. A region critical for IRES activity is distal from the translation start codon, bordering a region of conserved structured RNA as well as conserved unpaired sequence. This critical portion agrees with a previous report of the human insulin receptor 5’UTR. The D1D2 element can accommodate a variety of downstream RNA sequences and retain IRES activity. However, D1D2 is not independently sufficient even when folded correctly in cells. Sufficiency is only achieved when elements in the remainder of the UTR are replaced with generic linear linkers or TL hairpins, indicating sequence- and structure-dispensability of these regions. The requirement of these elements for sufficiency suggests a role for the remainder of the UTR which may be satisfied by highly divergent sequences among mammals. Depending on the mechanism of the Insr 5’UTR IRES, its function might be affected by nonproductive upstream initiation events or spatial constraints for start codon positioning. The dispensability of much of the UTR may hint at low structure requirements for cap-independent translation initiation.

## LIMITATIONS

Large RNA structures can present a challenge to secondary structure predictions. Though we selected a representative prediction, we acknowledge that our secondary structure model of the Insr 5’UTR requires refinement and corroboration. Even if our constraint data is found to be biologically representative by future studies, there may be structural features which our prediction tools cannot capture. When predicting RNA secondary structures from probing data, the probing may be influenced by RNA tertiary structure, and bases which appear unpaired by DMS reactivity may be involved in non-base-pairing interactions such as aromatic stacking.

## Supporting information

Table 1 Oligos Used

Table 3 Luciferase Assay Data

Table 2 Plasmid List

## DATA AVAILABILITY STATEMENT

Processed data (including raw and normalized DMS-MaPseq constraint files, RNA secondary structure prediction files, and statistical assessment of luciferase assay results) reported as the results of this study, as well as the plasmids used, are accessible through the corresponding author upon reasonable request. FASTQs used for DMS-MaPseq constrained RNA secondary structure prediction are deposited in the National Library of Medicine Sequencing Read Archive (SRA). Viral IRES DMS-MaPseq data accessions: SRR31191072, SRR31191071, SRR31191070, SRR31191069. Mouse Insr 5’UTR DMS-MaPseq data accessions: SRR31191090, SRR31191089, SRR31191080, SRR31191079, SRR31191078, SRR31191077, SRR31191076, SRR31191075, SRR31191074, SRR31191073, SRR31191088, SRR31191087, SRR31191086, SRR31191085, SRR31191084, SRR31191083, SRR31191082, and SRR31191081.

These accessions are separated by UTR variant and probing context. A current link to the DREEM analysis software can be found in Materials and Methods. Raw luciferase assay data is deposited in Table 3 in Supplementary Data.

## SUPPLEMENTARY DATA

Supplementary Data are available online.

## AUTHOR CONTRIBUTIONS

W.D., S.R., and M.T.M. conceived and designed the project. W.D. carried out all experiments with contribution from T.C.T.L. for initial DMS-MaPseq library preparation.

W.D. performed data analysis with input from S.R. and M.T.M. W.D. and M.T.M. interpreted the results and wrote the paper with input from S.R.

## ACKNOWLEDGEMENTS

We thank J.E. Haber for manuscript input and J.A. Doudna’s laboratory for the HCV IRES sequence.

