## Supplemental Figures for "Structure-informed mutagenesis identifies a conserved region critical for mouse insulin receptor 5’UTR IRES function"

**Supplemental Figure 1. Analysis pipeline and quality control metrics**

**(A)** The typical DMS-MaPseq analysis pipeline with an additional barcode splitting step added upstream to support the sequencing of many DMS-MaP Seq fragments at once. **(B-E)** DMS-MaPseq data on the HCV IRES in cells and *in vitro*. At left, representative scatterplots of mutation rates between two replicates in cells and *in vitro*, respectively. Red: As. Blue: Cs. Black: Gs and Us. At right, frequency histogram counting the number of mutations per 150x150 paired read. **(F-I)** The same DMS-MaPseq data for the EMCV IRES. **(J-M)** DMS-MaPseq data for Insr probed in tandem with HCV and EMCV. These data were used for secondary structure predictions.

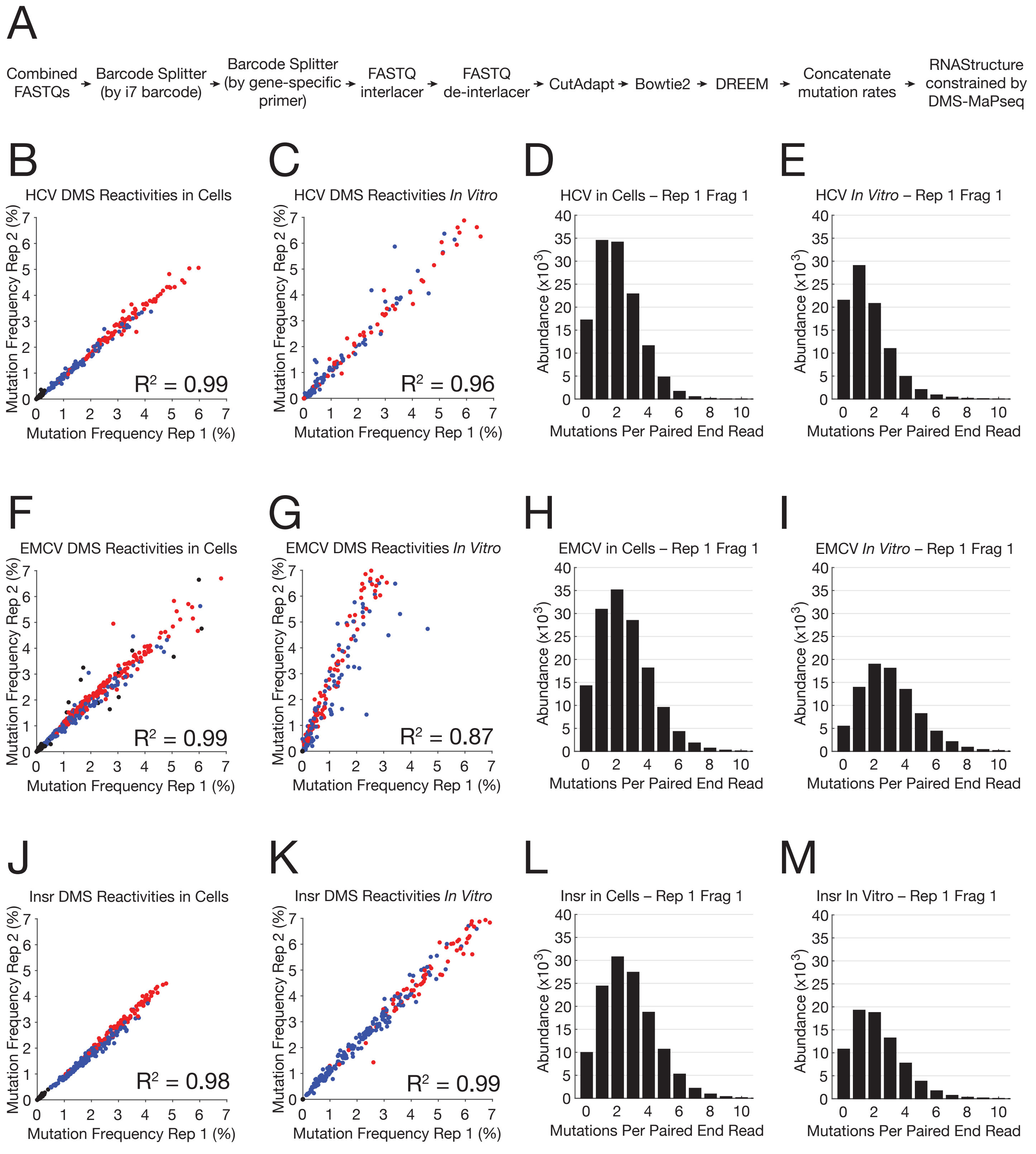

**Supplemental Figure 2. Known structures of HCV and EMCV viral IRESes used for comparison**
**(A)** The known structure of the HCV IRES, adapted from Kieft et al. 1999. This isolate differs by 4nt from the probed structure, annotated in the figure. In cell DMS-MaPseq data is superimposed for comparison. AUROC between this structure and both in-cell and *in vitro* DMS-MaPseq data is indicated. **(B)** The known structure of the EMCV IRES, adapted from Pilipenko et al. 1989. The probed structure includes adjacent domains. Indexes are based on the probed structure. AUROC comparisons of this known structure to *in vitro* and in cell DMS-MaPseq data are included. In cell DMS-MaPseq data is superimposed.

# A

**HCV Known Structure (*in vitro*)**
Source: Kieft et al. 1999
In cell DMS-MaPseq data superimposed.
AUROC = 0.76 vs. in cell DMS-MaPseq
AUROC = 0.80 vs. *in vitro* DMS-MaPseq

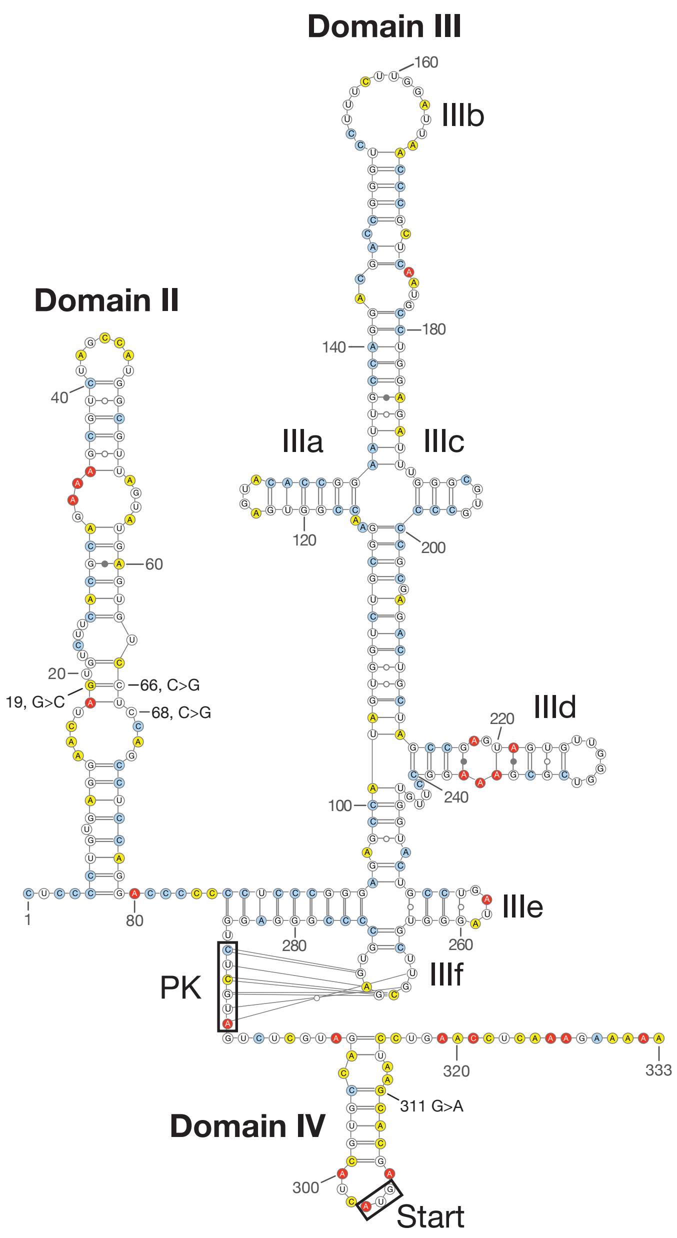


# B

**EMCV Known Structure (*in vitro*)**
Source: Pilipenko et al. 1989
In cell DMS-MaPseq data superimposed.
AUROC = 0.80 vs. in cell DMS-MaPseq
AUROC = 0.81 vs. *in vitro* DMS-MaPseq

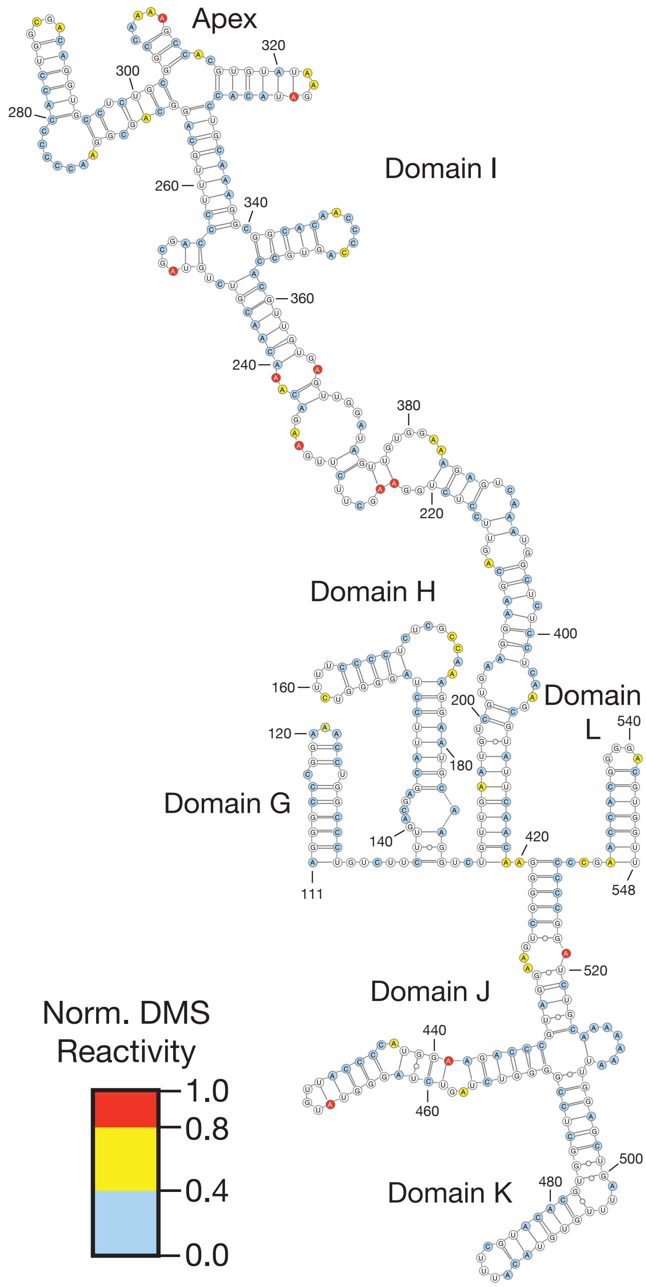

**Supplemental Figure 3. Partitioning and conservation of the Insr 5'UTR**

**(A)** The Insr 5'UTR prediction from Figure 4b divided into domains with indexes indicated. (B) Full tracks of Insr 5'UTR conservation among mammals. Green peaks: conservation among 30 mammals. Magnitude letters: conservation among 470 mammals. vTop: human. Bottom: mouse. Green peaks indicate conservation among 30 mammals. Magnitude letters indicate conservation among 470 mammals, with hats indicating fixed nucleotides. RNA structure domains discussed in this report are annotated.

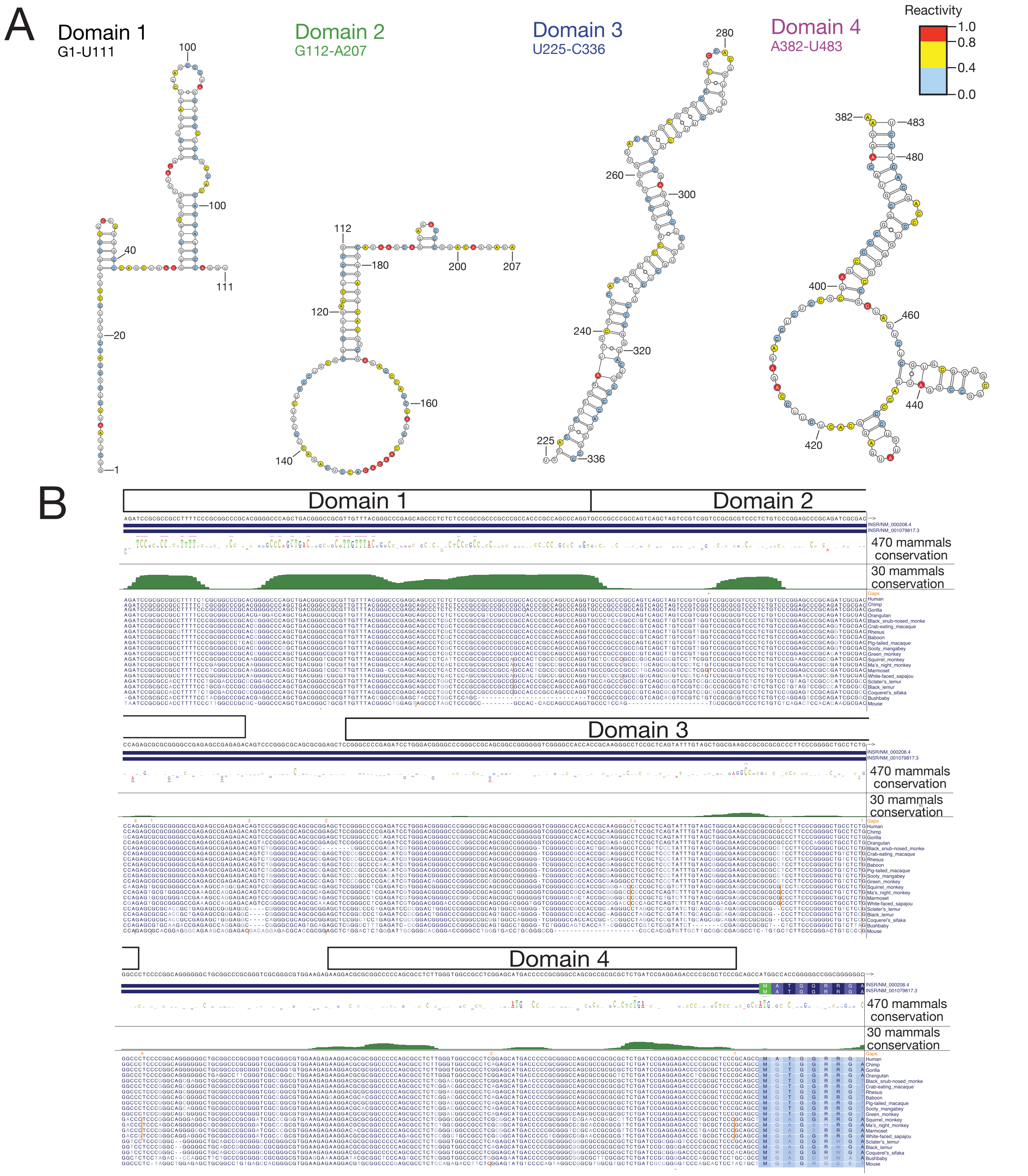

**Supplemental Figure 4. All tetraloop hairpin mutants assayed**

**(A)** Bicistronic luciferase assay results from tetraloop hairpins walking down each of the predicted domains. Statistics: Tukey's HSD of each mutant compared to the full length Insr 5'UTR *** = $p < 0.0005$, ** = $p < 0.005$, * = $p < 0.05$. Not significant: ns. Location of mutations is depicted. **(B)** Cartoon of tetraloop hairpins' locations with positions noted.

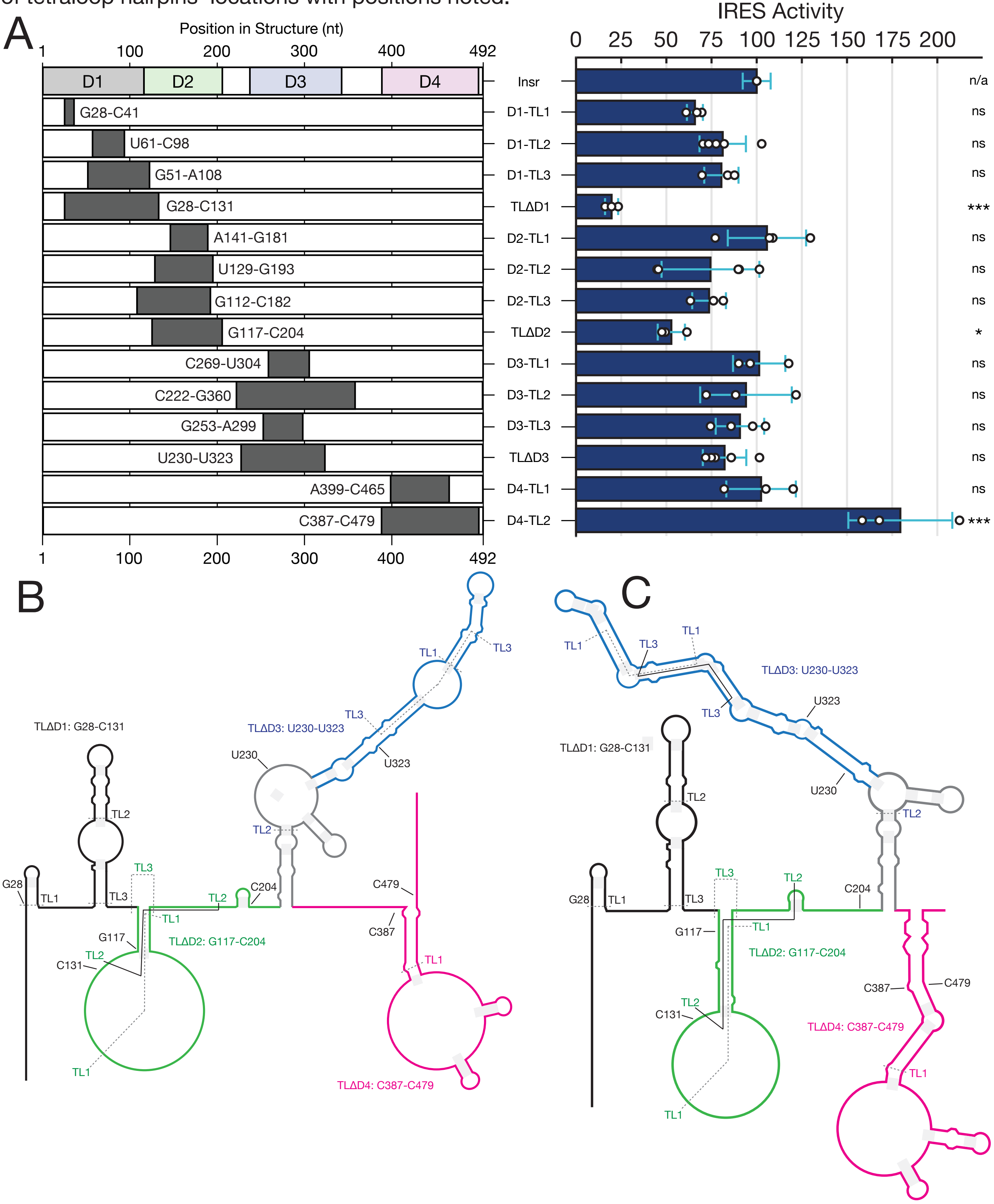

**Supplemental Figure 5. DMS-MaPseq on mutants *in vitro***
**(A-F)** Coefficient of determination on As and Cs plotted in windows as described. Mutations' position and nature (Δ = deletion, TL = tetraloop hairpin replacement) are indicated.

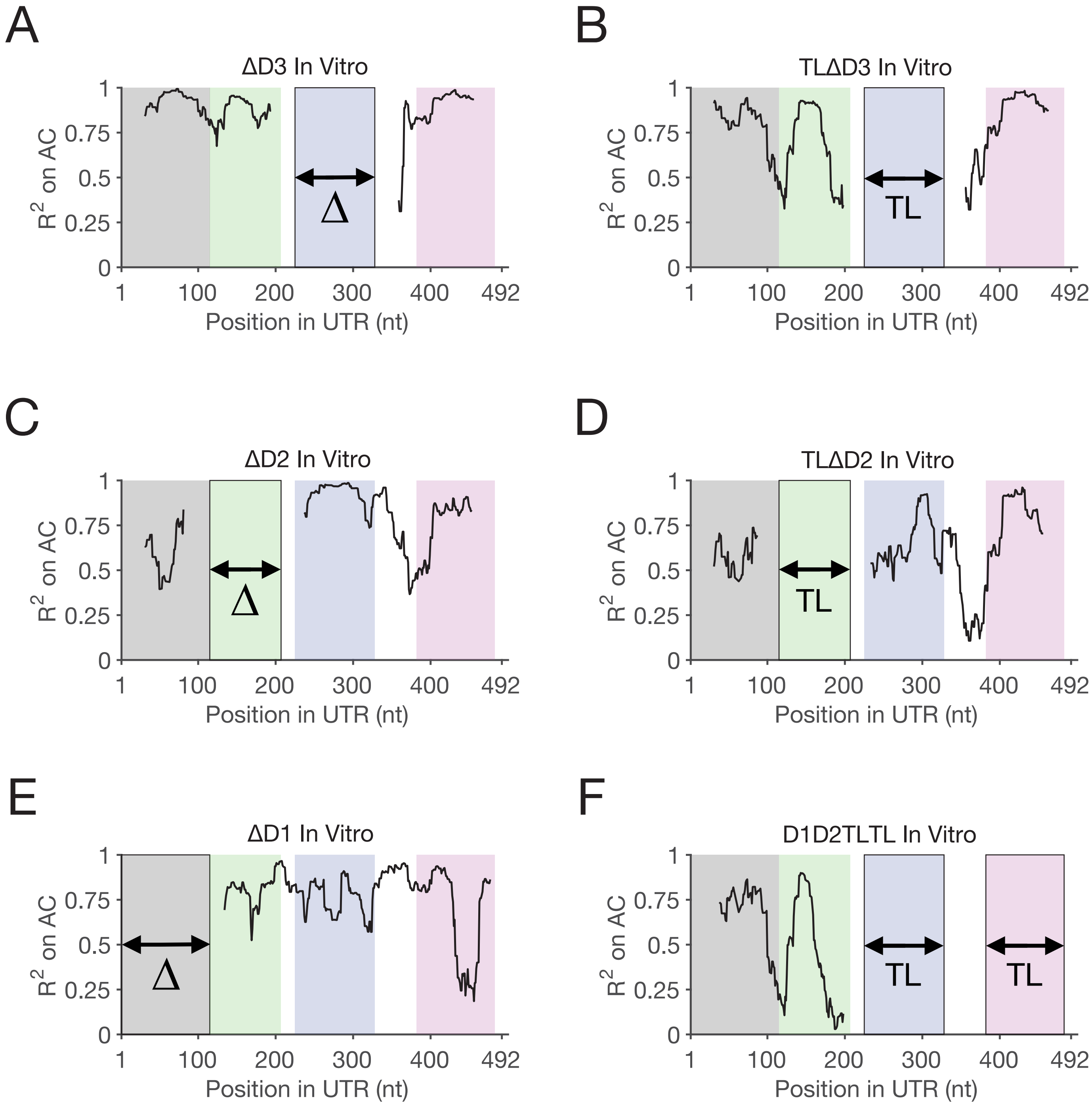

**Supplemental Figure 6. Performance of disruption and solo stem mutants**
**(A)** Bicistronic reporter results for sequence disruptions (detailed in Table 1) with statistics as previously described. **(B)** Assay results of each domain's individual sufficiency for IRES with the rest of the UTR removed.

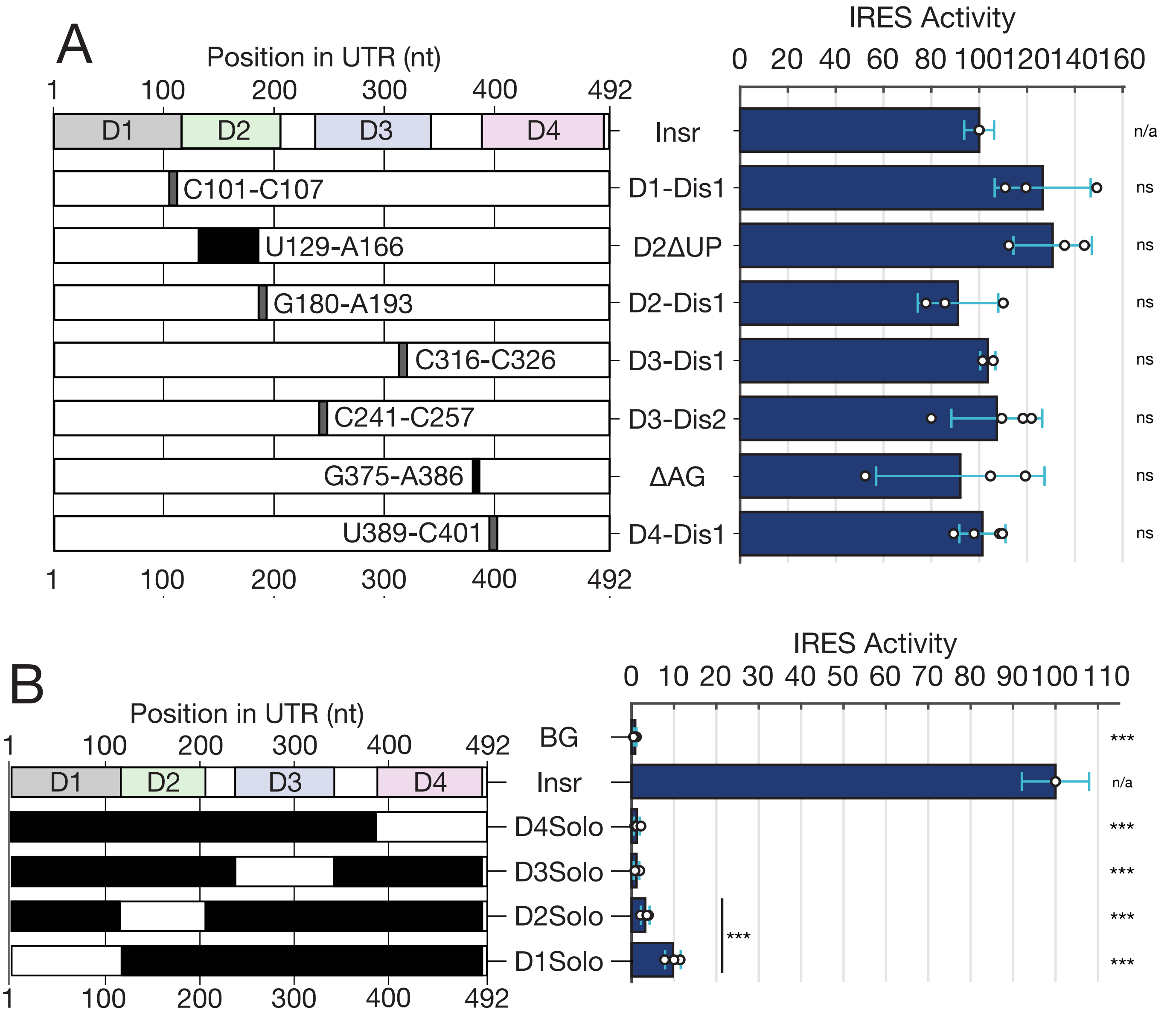
